# Towards a scalable effective connectivity estimation in fMRI

**DOI:** 10.64898/2026.10.26.761780

**Authors:** Benedetta Mariani, Luca Taffarello, Giorgia Baron, Giacomo Barzon, Federico Calesella, Matthieu Gilson, Stefano De Marchi, Alessandro Chiuso, Antonino Vallesi, Samir Suweis, Marco Zorzi, Alessandra Bertoldo, Michele Allegra

## Abstract

Characterizing directed connections between brain regions is fundamental for understanding the brain’s network organization and how it supports cognitive processes. However, a reliable methodology allowing assessment of effective connectivity (EC) at the whole-brain level and across high-resolution parcellations has remained elusive. This limitation reflects a trade-off between model detail and scalability. Mechanistic approaches such as Dynamic Causal Modeling (DCM) provide a principled description of directed interactions but become impractical for large parcellations, whereas scalable methods based on linear-system inversion often fail to reliably recover key features of the underlying effective connectivity. Here, using data from the Human Connectome Project S1200 release, we show that the multivariate Ornstein-Uhlenbeck (mOU) approach for EC estimation can be adapted to yield EC estimates that are in broad agreement with those of an advanced DCM, at least when the spatial variability of the hemodynamic response falls within the physiological range characteristic of the healthy adult brain. Furthermore, thanks to its low computational cost, mOU can be easily scaled up to large neuroimaging cohorts and high-resolution parcellations. High-resolution, mOU-based EC outperforms functional connectivity for single-subject connectome finger-printing. In addition, it enables a high-resolution characterization of the brain’s hierarchical organization and modular architecture by identifying directed interactions within and between communities and revealing driver and receiver regions. Overall, our methodology provides a reliable and scalable EC estimation pipeline of wide applicability to cognitive and network neuroscience.

## Introduction

Cognitive processes are typically not localized, but require the interplay of several brain areas [1, 2, 3]. Common metrics to characterize large-scale brain networks - structural connectivity (SC), which quantifies physical axonal connections between brain regions through diffusion tensor imaging or tract-tracing, and functional connectivity (FC), which measures statistical correlation in functional magnetic resonance imaging - are typically non-directional. Therefore, although they provide valuable insight into the large-scale organization of the brain in health and pathology [4, 5, 6, 7], they cannot be employed to reconstruct directional relations between brain regions, for which information about the directionality or asymmetry of interaction is essential. Effective connectivity (EC) characterizes the directed causal influence one neural population exerts over another [8], which depends on both synaptic strength (e.g., types and concentrations of neurotransmitters) and dynamic properties such as neural excitability [9]. EC is essential for understanding information flow and the propagation of signals across the brain, including bottom-up sensory stimuli, top-down feedback, and external neuromodulation [10, 11, 12, 13, 14, 15, 16, 17, 18, 19, 20, 21, 22, 23, 24, 25]. Moreover, EC uniquely enables the characterization of directed hierarchies by identifying asymmetric influences between regions and distinguishing source, relay, and sink roles within the network. However, contrary to FC and SC, EC is based on a parametric generative model estimation rather than directly inferred from data.

The most common approach for estimating effective connectivity in fMRI is Dynamic Causal Modeling (DCM) [26], which employs biophysically informed generative models and Bayesian inference to explain observed data in terms of hidden neural states and directed interactions. DCM relies on a two-layer model, including a dynamical model at the level of neural activity, and an observation model encapsulating region-specific hemodynamic response functions (HRFs), mapping neural activity into the experimentally measured blood-oxygen-level-dependent (BOLD) signals [27]. Due to the nonlinearity of the observation component, model inversion is computationally demanding. The original, deterministic formulation of DCM [9] was conceived for task fMRI data and was generally restricted to small networks (*∼*10 regions). A stochastic DCM, driven by endogenous random fluctuations, was later developed to describe the resting state [28], and then simplified in [29] by postulating a linear model for haemodynamic responses. This step considerably facilitated model inversion, enabling the application of the model to networks of up to 36 regions. Further progress came from the work in [30], where the authors included a sparsity-inducing prior and a statistical linearization of the haemodynamic response. Despite these simplifying adjustments, the resulting sparse DCM maintains a heavy computational load, preventing its application to networks with *>* 90 nodes [31]. These limits are not present in regression DCM [32, 33], which can infer effective connectivity among *>* 100 brain regions, but at a huge price: resorting to a coarse hemodynamic model based on a known, region-unspecific hemodynamic response.

An alternative framework to DCMs is the multivariate Ornstein–Uhlenbeck (mOU) model proposed by Gilson et al. [34, 14]. Like stochastic DCMs, mOU assumes that whole-brain activity follows linear stochastic dynamics. Unlike DCM, however, it does not include an observation model, sacrificing the ambition to model a region-specific hemodynamic response - a simplification that neglects inter-regional time shifts in HRFs, as in regression-DCM [32]). Moreover, mOU estimates directed connectivity by matching empirical covariance structures through convex optimization, instead of relying on likelihood-maximization techniques. Thanks to its simplifying assumptions, mOU is highly computationally efficient, and it can be applied to large-scale resting-state datasets. However, it remains unclear whether this gain in scalability comes at the cost of reliable EC inference. In particular, the consistency of mOU-based EC estimates with those obtained from DCM, under substantially richer modeling assumptions, and their robustness as network size increases have not been systematically assessed. Here, we address these questions by benchmarking mOU against DCM and systematically testing its ability to recover effective connectivity across increasingly large brain networks.

Here, we use a large-scale fMRI data set of healthy participants from the Human Connectome Project [35] (*N* = 1013) to systematically test the consistency of mOU-based EC estimates (i) with sparse DCM-based EC estimates, (ii) across parcellations with different degrees of spatial resolution, and (iii) across different recordings sessions of the same participants. The choice to compare mOU with sparse DCM, as opposed to other models in the DCM class, is due to the fact that sparse DCM is currently the only DCM representative allowing EC estimation from resting-state fMRI at the whole-brain level (regression DCM essentially skips the observation model, like mOU, thus losing the most characteristic trait of DCM).

Our manuscript is organized as follows. Firstly, we show that the original estimate of the self-decay timescale (*τ*) in mOU is not robust under standard BOLD preprocessing. We therefore introduce an alternative estimate of *τ*_*\**_. We show that *τ*_*\**_ lies within a broad range of *τ* values yielding stable algorithmic convergence, plausible EC asymmetry, and maximal consistency with sparse DCM estimates. Secondly, we address the scalability of mOU EC estimation by applying it to Schaefer parcellations at increasing resolutions (62, 100, and 200 parcels). While sparse DCM is restricted to the lowest resolution because of its computational cost, mOU can be efficiently applied to substantially larger networks. We assess the consistency of high-resolution estimates by coarse-graining them to the 62-ROI scale and comparing them with EC inferred directly at that resolution. Thirdly, we assess the reliability of mOU-EC estimates at the single-subject level through subject fingerprinting, testing whether EC patterns can identify individuals across repeated resting-state fMRI sessions and comparing their performance with FC. Finally, we exploit the directed nature of EC to characterize brain-network organization using measures unavailable to conventional FC, including source-sink properties and directional hierarchies between brain regions.

Overall, our work demonstrates that mOU-based EC can be taken as a reliable, scalable proxy of DCM-based EC, opening the way to high-resolution analyses of directed connectivity and hierarchical relations in the brain.

## Results

We analyzed resting-state fMRI data of healthy participants from the Human Connectome Project (HCP) 1200-subject release [35]. We used the left-right (LR) runs from the first session (REST 1) for our main analysis, and included LR runs from the second session (REST 2) to perform fingerprinting. Subjects were filtered on the basis of the availability of both REST 1 and REST 2 runs, which resulted in a final number of 1013 subjects. We fitted the mOU model at the individual participant level, using the 62 ROIs parcellation derived from the Schaefer-100, Schaefer-100 and Schaefer-200 parcellations (see Methods). Throughout, *Q*_0_ and *Q*_lag_ denote theoretical covariances at zero lag and lag *t*_lag_, while 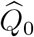 and 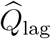 denote their empirical estimates; 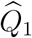 denotes the empirical covariance at a lag of one TR. Hats on *J* and Σ likewise denote fitted parameters.

### Timescale calibration in the mOU model

Resting-state BOLD time series are typically low-pass filtered, as this step removes high-frequency cardiac and respiratory artifacts and retains signal related to major resting-state networks [36, 37]. Low-pass filtering at 0.1 is the standard for functional connectivity analysis, and it was also used in many works implementing whole-brain DCM [38, 39, 40]. In fact, this step significantly improves within-subject consistency of EC estimates (as shown below). However, current implementations of the mOU model [14, 41] struggle when applied to low-pass-filtered fMRI data. The mOU process is defined as

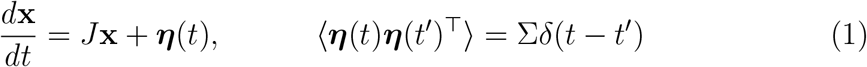

where the Jacobian is defined as 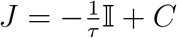, *C* being the effective connectivity (EC) matrix and I the identity matrix, and ***η***(*t*) is a Wiener process with covariance matrix Σ (see Methods for details). The parameter *τ*, which determines the self-decay rate of the model and acts as a global time scale for the mOU process, was estimated by Gilson et al. [14, 42] by fitting an exponential only to the first two points of the autocorrelation function. For lowpass-filtered data, as shown in Fig. 1(g), this choice leads to a gross overestimation of the time scale *τ* (purple). This issue cannot be simply solved by including more points in the estimation of the autocorrelation function: for example, by including the third point (orange in Fig. 1(g)), we only mildly alleviate the overestimation. The problem is that the autocorrelation function of time-filtered data does not exhibit the predicted exponential decay.

**Figure 1:**
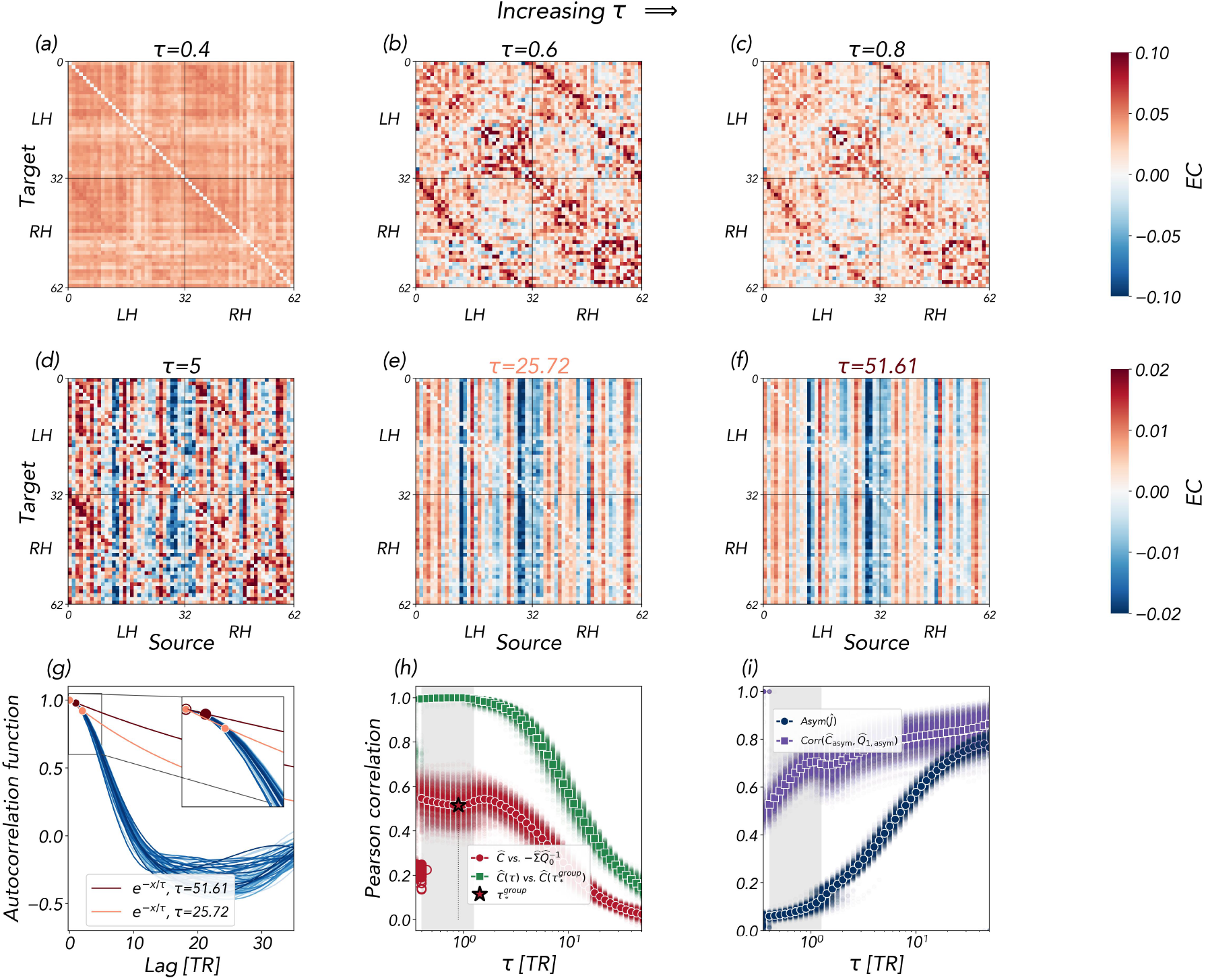
Dependence of effective connectivity on the decay timescale. **(a–f)** Effective connectivity (EC) matrices estimated for a representative participant at increasing values of *τ* . Panels (e,f) correspond to the decay timescales illustrated in (g). **(g)** Regional autocorrelation functions and exponential fits using zero lag and either one or two TRs. The inset shows the short-lag behavior. **(h)** Pearson correlation between EC and the noise-weighted precision matrix 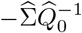 (red), and between EC (*Ĉ*) at each *τ* and EC at the common reference timescale 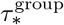 (green). The star marks 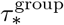, selected as the minimum of the mean red curve within the convergence range. Open red circles indicate initial outliers of the red curve excluded from timescale selection. All correlations are computed among off-diagonal entries. **(i)** Jacobian asymmetry (blue) and Pearson correlation between the antisymmetric components of *Ĉ* and the empirical lag-one covariance 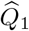(purple). In **(h**,**i)**, transparent points represent individual participants (*n* = 1013) and solid curves indicate their means. Gray shading indicates timescales satisfying the convergence for at least 70% of participants.

We systematically analyzed results obtained when fixing different values of *τ* (measured in units of the repetition time *TR* = 0.72*s*) and retrieving the EC matrix *C* with the standard Lyapunov-optimization procedure (Methods). In Fig. 1(a-f), we show EC matrices obtained for a representative HCP subject. For *τ ≲* 0.4, the EC was highly symmetric and lacked directionality (figure 1(a), and highly correlated (*r*=0.9) with the covariance matrix 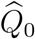). For other subjects, small values of *τ* often led to an even poorer outcome, namely, convergence of the EC estimation to either a null matrix or NaNs. On the other hand, for *τ ≳* 10 the EC was highly asymmetric (Fig. 1(d, e, f)), featuring the emergence of entire columns with consistently larger-than-average entries, depicting a biologically implausible scenario where a few nodes disproportionately affect all the rest of the network. However, in a relatively broad interval of intermediate values of *τ*, EC matrices were consistent and biologically plausible (Fig. 1 (b,c)). Typically, we obtained stable solutions only for a relatively wide interval of *τ* values, denoted here as “convergence range”. Outside this interval, the loss function continued to decrease, failing to reach a plateau even when the maximum number of iterations was increased, indicating that the inferred EC had not reached a stable optimum. For most subjects, the convergence range was [0.5,1.3] (see Fig. S1), while *τ >* 2 always fell outside the convergence range, and, as mentioned, *τ* ≤ 0.4 often led to a null matrix or NaN. These findings suggest that suitable solutions are found at intermediate values of *τ* . In Fig. 1(h,i), gray shading identifies timescales satisfying the convergence in at least 70% of participants. According to the Lyapunov equation, *J* should admit the following decomposition [34, 43]:

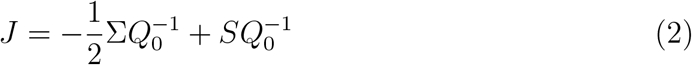

Here *Q*_0_ denotes the stationary model covariance; empirical comparisons use the sample covariance 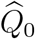. The first term, which is symmetric and equal to the precision matrix 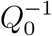 if Σ is proportional to the identity, represents the time-reversible part of the dynamics, which obeys detailed balance. The second term is linked to the differential cross-covariance 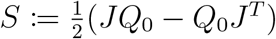 [43, 40] (the antisymmetric component of the differential covariance *JQ*_0_), which represents the time-irreversible component of the dynamics, namely the pairwise directional imbalance that breaks detailed balance [43].

In Fig. 1(h), we examine the off-diagonal Pearson correlation between the inferred EC and the noise-weighted precision reference, 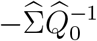(red). Throughout the intermediate range of *τ* associated with stable convergence, this correlation remained high, suggesting that the reversible component contributes substantially to the spatial pattern of inferred connectivity. Within this range, the correlation exhibited a local minimum, suggesting that there is a specific point (*τ*_*\**_) where inferred connectivity departs most strongly from the reversible reference, potentially leaving greater scope for irreversible contributions. In Fig. 1(i), we analyze the asymmetry of the inferred Jacobian, measured as Asym 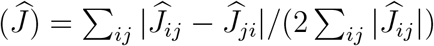. The asymmetry (blue curve) increased monotonically with *τ*, slowly in the convergence range and wildly beyond, indicating a contribution of non-reciprocal coupling that remains stable only within the convergence range. We also analyzed the correlation between the antisymmetric components of the estimated Jacobian and the empirical lag-one covariance 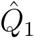, which characterizes the link between the directional pattern inferred by the model and the temporal asymmetry in the data. This correlation (purple curve) reached a peak at *τ*_*\**_ (Fig. 1i), suggesting that directionality in EC links is maximally aligned with temporal asymmetries in the data.

In summary, we identified an intermediate-*τ* regime in which mOU estimation converged and connectivity patterns were consistent. We selected a common reference timescale 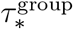 as the minimum of the group-averaged red curve within the convergence range (excluding the trivial points found at low *τ*). It is worth remarking that there is no need for a fine-tuned selection of *τ* : within the convergence range, estimated EC matrices always remained highly correlated (*r >* 0.99) with EC matrices computed at the common reference timescale (Fig. 1h, green).

### Comparison between mOU and Sparse DCM

We compared EC estimates provided by mOU (where *τ* was fixed as explained in the previous section) with those provided by sparse DCM. To this aim, we restricted our attention to the off-diagonal entries of the Jacobian matrix *J*, as the mOU model (contrary to sparse DCM) does not provide region-specific estimates of the diagonal elements (i.e., self-decay constants). For this analysis, we used a coarse-grained version of the Schaefer parcellation comprising 62 parcels (obtained from the standard 100-parcel Schaefer parcellation by aggregating some regions, as detailed in the Methods section). Moreover, we restricted the analysis to a subset of 144 subjects already analyzed in Ref. [40]. In fact, given the heavy computational load of DCM, it would have been computationally demanding to work with the entire cohort of *∼*1000 subjects, highlighting the difficulty of employing sparse DCM in the analysis of large-scale neuroimaging databases with one thousand subjects or more. Note that mOU vastly outperformed DCM in terms of computational cost (see Methods for the typical cost of the two approaches). As shown in Fig. 2, we found a good agreement between the two methods. Panels (a-f) show subject-level results for the two subjects with the best and worst alignment between mOU and DCM (in terms of correlation between the respective EC matrices). Panels (g-i) show group-level (average) results. To quantify the alignment between mOU and DCM, we computed Pearson’s correlation between the respective EC matrices (matrices were vectorized, and the diagonal was removed). The insets in Fig. 2(c) and (f) show the distribution of correlation values over all subjects, with arrows highlighting the location of the correlation value observed in the main plots. Correlation values are high for all subjects (mean=0.82; 5th percentile=0.77; 95th percentile=0.86). Fig. 2(i) shows a scatter plot between the entries of the group-level EC matrices obtained with mOU and sparse DCM: the group-level correlation is very high, *R* = 0.89. We note that for positive links, the correlation would be even higher, while negative links are less correlated.

**Figure 2:**
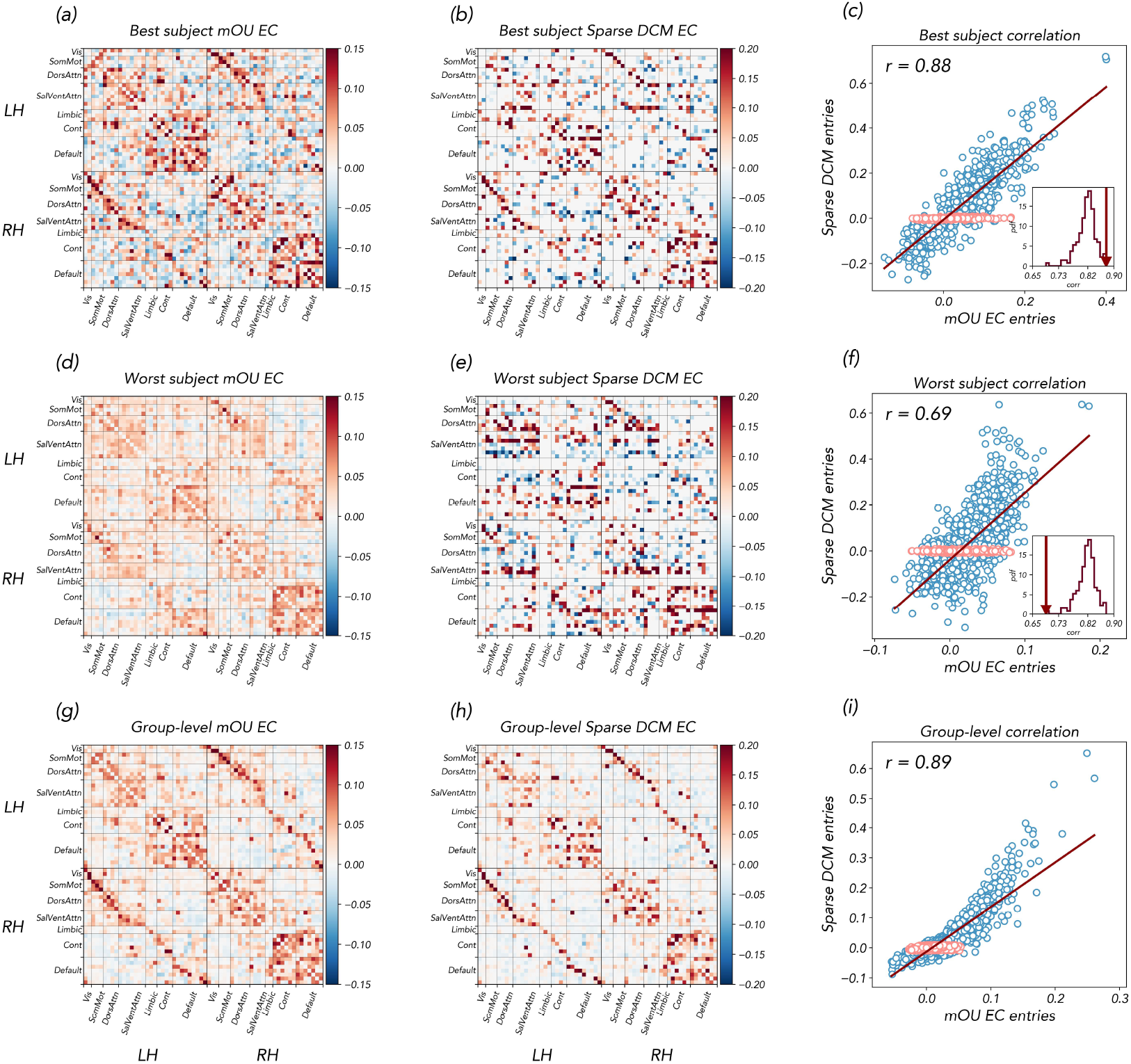
Comparison between mOU- and sparse DCM-based effective connectivity estimates. **(a–c)** Effective connectivity matrices estimated with the mOU model and sparse DCM for the subject showing the highest correspondence between the two methods, together with the edge-wise relationship between the corresponding non-zero connectivity estimates. Entries classified as absent by sparse DCM are shown in the scatter plot in pink. The inset reports the distribution of subject-level correlations across the full sample, with the selected subject indicated by the vertical arrow. **(d–f)** Same comparison for the subject showing the lowest correspondence between mOU and sparse DCM estimates. **(g–i)** Group-average mOU and sparse DCM effective connectivity matrices and their edge-wise association. For all scatter plots, the solid line indicates the linear regression fitted to the non-zero sparse DCM entries, and the Pearson correlation coefficient is reported in the upper-left corner.

Thus, despite their different assumptions (the inclusion of a sparsity prior and a region-dependent haemodynamic response in DCM, and the different inference procedures), the agreement between the two methods is remarkably good. This result was partly surprising because inter-regional differences in haemodynamic response functions (HRFs) would be expected to cause noticeable discrepancies between mOU and DCM. Many features of the HRF may affect EC estimates, including amplitude, undershoot and latency. The latter could be particularly relevant: a difference Δ*ℓ*_*ij*_ = *l*_*i*_ − *l*_*j*_ *>* 0 between the latencies *ℓ*_*i*_ and *ℓ*_*j*_ of regions *i* and *j* would be expected to cause a systematic lag in the corresponding BOLD time series, which mOU would mis-attribute to the presence of an EC link *C*_*ij*_ between the leading region (*j*) and the lagging region (*i*). We therefore hypothesized that the two methods for estimating effective connectivity are in reasonable agreement when the spatial variability of the hemodynamic response falls within the physiological range characteristic of the healthy adult brain. Indeed, we found only moderate differences in HRF latencies across regions – whose values are in the range [3.6 s, 6.4 s] (5th and 95th percentile of the latencies including all the subjects and all the regions) (see supplementary Fig. S2) We found that these differences in HRF latencies had only a moderate effect on the estimation difference between 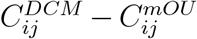. In fact, as Fig. S3 shows, the correlation between group-level Δ*C*_*ij*_ and group-level lag difference Δ*ℓ*_*ij*_ across different links is low, and often the biggest differences between the two methods are not necessarily due to a latency difference among the regions (see Fig. S3c). On the other hand, we found that for some links, there is a significant effect (significant negative correlation) between Δ*C*_*ij*_ and Δ*ℓ*_*ij*_ across subjects (Fig. S3e) (*R* = −0.76 for the most significant link): this means that, for some links, when a subject presented a latency difference among the related regions, mOU tended to overestimate the strength of the EC link from the leading to the lagging region. This hints at the possibility of this estimation bias (even though, as mentioned, we did not observe a systematic correlation across links between Δ*C*_*ij*_ and Δ*ℓ*_*ij*_), and warns against the blind application of the mOU method in situations in which HRF differences might be more marked, for example in pathological conditions.

### Scalability of the mOU model

To test the scalability of mOU to larger parcellations, we ran the model on different versions of the Schaefer atlases, going from coarser (62 parcels) to finer ones (100, up to 200 parcels; see Methods for details on the mapping between different parcellations). mOU did not encounter any computational difficulty in EC matrices of size 100 and 200 (Fig. 3, top row)(contrary to DCM, for which model inversion would be computationally unfeasible even with 100 nodes). Example EC matrices for different parcellations are shown in Fig. 3, top row. We tested the consistency of estimates at different resolution levels by averaging EC values over all high-resolution parcels belonging to the same low-resolution parcel [42] (see Methods for details). In this way, we obtained low-resolution EC matrices starting from high-resolution ones (instead of running the model directly at a low resolution). Results of this coarse-graining step are shown in Fig. 3, top row. Importantly, we obtained a noteworthy agreement between the 62-parcel EC matrices obtained by coarse-graining either the 100- or the 200-matrices, and those obtained directly by running mOU on the 62-parcel time courses. The former are shown in Fig. 3, middle row. In the bottom row of Fig. 3, we also show the entry-wise alignment between the low-resolution EC matrices obtained directly and those obtained with the coarse-graining procedure. To quantify the alignment, we computed Pearson’s correlations: we obtained *R >* 0.9 for both cases. While Fig. 3 shows group-level ECs, a similar picture (and similar values of the correlation) is obtained for individual EC matrices. In summary, although the number of parameters increases, the mOU inversion is computationally tractable, and the resulting mou-EC estimates are consistent with mou-EC estimates obtained with low-resolution parcellations.

**Figure 3:**
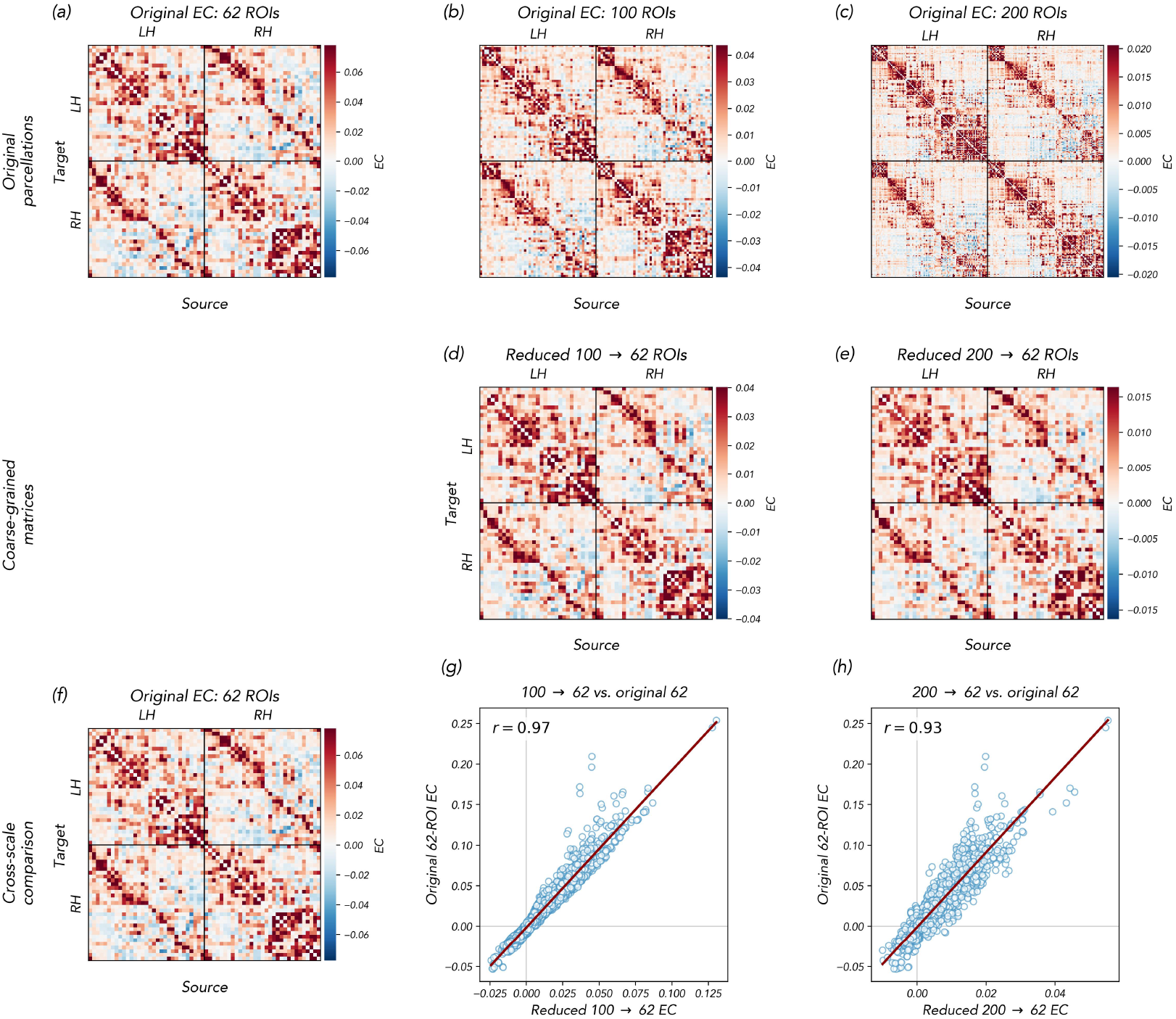
Consistency of mOU-effective connectivity estimates across parcellation scales. **(a–c)** Group-average mOU-effective connectivity matrices estimated using parcellations with 62, 100, and 200 regions of interest, respectively. The black lines separate the left and right hemispheres. **(d–e)** Effective connectivity matrices obtained by coarse-graining the 100- and 200-region estimates to the 62-region parcellation. **(f-g-h)** Edge-wise comparison between the original 62-region mou-EC estimate and the corresponding coarse-grained matrices derived from mou-EC estimates obtained with the 100- and 200-region parcellations. Solid lines indicate linear regressions, and Pearson correlation coefficients are reported within each panel.

### Single-subject consistency and fingerprinting

To further assess the reliability of mOU-derived effective connectivity (EC) at the individual level, we evaluated whether EC matrices remain sufficiently stable across repeated resting-state acquisitions to support subject identification. Following the terminology introduced by Finn et al. [7], this procedure is commonly referred to as *connectome fingerprinting*. In this framework, identification is considered successful when the connectivity matrix of a subject obtained in one scanning session is more similar to that subject’s matrix from a second session than to the matrices of all other individuals in the cohort. For our purpose of evaluating reproducibility and consistency, we employed the first proposed method to study fingerprinting ([7]), i. e., simply computing the Pearson correlation between each REST 1 and each REST 2 scan. We obtain thus an identifiability matrix, whose diagonal elements are called *self-identifiability*, while the off-diagonal ones *others-identifiability* [44]. Fingerprinting is considered successful if the self-identifiability of a subject is higher with respect to their others-identifiability. Previous fingerprinting studies focused primarily on functional connectivity (FC), showing that FC captures stable subject-specific features of large-scale brain organization [7, 44, 45]. However, most of these studies considered relatively small cohorts (*∼*100 subjects) and high-resolution parcellations (typically 268 or 360 regions) [7, 44], conditions that favor subject identifiability. More recently, fingerprinting analyses have been extended to larger cohorts from the HCP dataset [46]. Here, we used fingerprinting as a benchmark for evaluating the reproducibility and subject-specificity of EC estimates. In particular, we compared the ability of FC and mOU-derived EC to identify individuals across the two resting-state sessions available in the HCP dataset. We tested fingerprinting for different cohort sizes (by randomly subsampling the whole cohort to obtain smaller groups) and different parcellation resolutions. We found that FC-based identification strongly depends on the spatial resolution of the parcellation. For relatively coarse parcellations (62–100 ROIs), FC fingerprinting accuracy reached at most *∼* 70% when fewer than 100 subjects were considered but decreased substantially for larger cohorts (Fig. 4). This reduction reflects the fact that, although FC matrices of the same participant are highly reproducible across sessions, in a large cohort, it is more likely to find another individual’s FC matrix also having high similarity with any given FC matrix(Fig. 4c). As expected from previous studies [7, 44], FC accuracy improved considerably when finer parcellations were employed (Fig. 4a), indicating that FC fingerprinting critically relies on high spatial resolution to capture sufficiently individualized patterns of connectivity.

**Figure 4:**
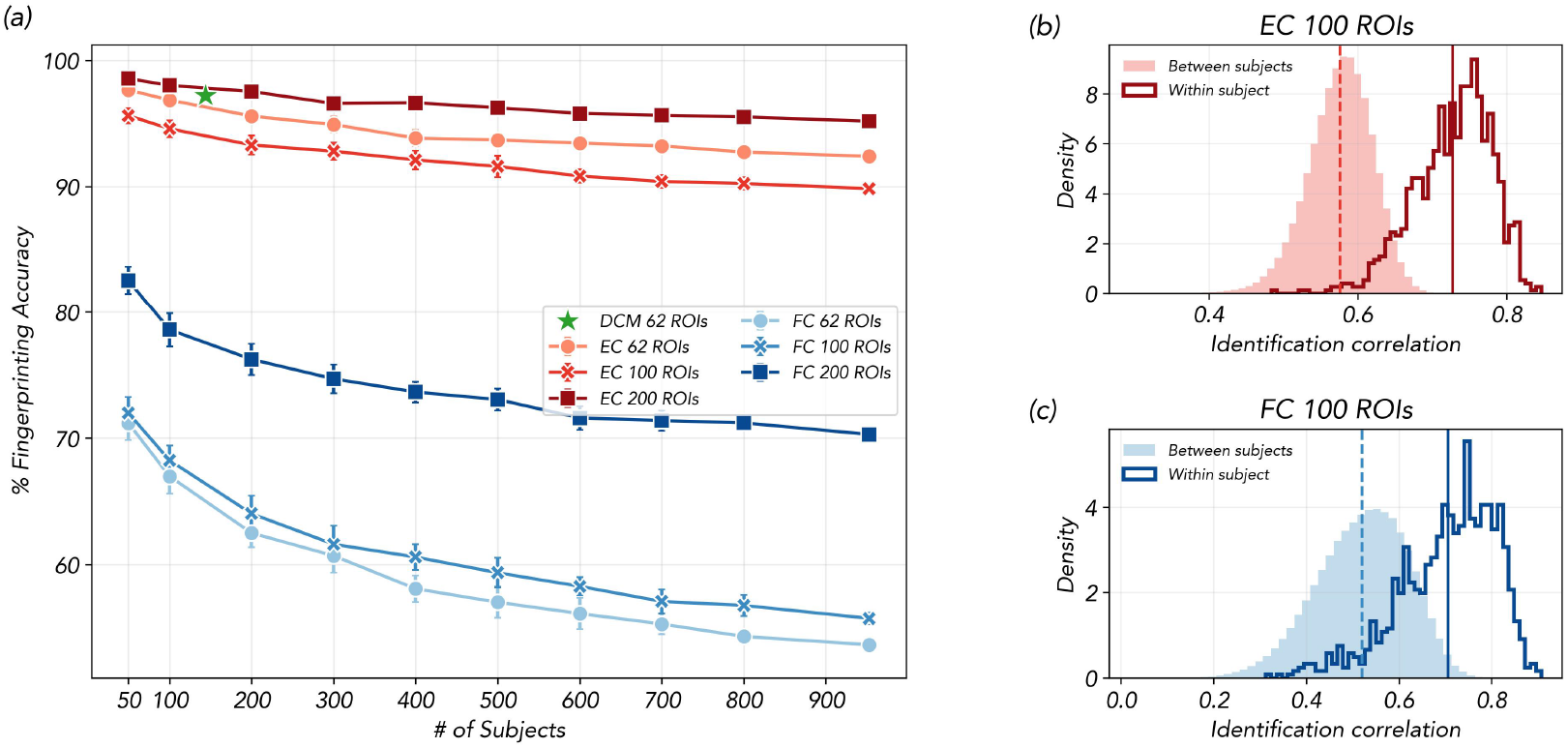
Fingerprinting performance of effective and functional connectivity across parcellation scales. **(a)** Identification accuracy between REST 1 and REST 2 as a function of the number of subjects for effective connectivity (EC) and functional connectivity (FC), evaluated using parcellations with 62, 100, and 200 regions of interest. Error bars indicate variability across repeated subsampling realizations. The green star indicates DCM fingerprinting accuracy using 62 ROIs (95.24%). **(b–c)** Distributions of within-subject correlations, corresponding to the diagonal elements of the identification matrix between REST 1 and REST 2, and between-subject correlations, corresponding to its off-diagonal elements, for EC and FC, respectively. A larger separation between the two distributions indicates stronger subject-specific identifiability.

In contrast, mOU-derived EC matrices exhibited both strong within-subject consistency across sessions and increased inter-subject variability (Fig. 4b). As a consequence, EC-based identification accuracy remained consistently high across all experimental conditions. In particular, accuracy was always greater than 85%, even when considering the largest cohort (*N* = 1013) and the coarsest parcellation (62 ROIs). Increasing the spatial resolution further improved performance, although the dependence on parcellation granularity was substantially weaker than in the FC case. Similar accuracy performances are observed with the sparse DCM EC (highlighed by the green star in Fig. 4a), with the only limitation that the analysis is limited to 144 subjects, and to a single parcellation (62 ROIs, as in the previous sections). Fig S6 reports the identifiability distributions for the DCM matrices.

As a control, we repeated the same fingerprinting analysis on mOU-ECs derived from non-filtered data (Figure S5). This study confirmed that excluding the temporal filtering step (the 0.01-0.1 Hz band-pass) reduces the self-identifiability of EC across REST1 and REST2 sessions. Moreover, in the non-filtered data case, fingerprinting performance was substantially reduced, with identification accuracy dropping to approximately 50%, when considering the full cohort.

These results suggest that directed interactions inferred through mOU capture subject-specific dynamical features that remain stable across repeated resting-state acquisitions while also preserving discriminability between individuals. Notably, this advantage with respect to FC is not simply explained by the larger number of independent entries in EC compared with the symmetric FC matrix. Indeed, as shown in Fig. S4, a similar trend is observed when fingerprinting is performed using symmetrized EC matrices. This suggests that mOU-derived EC captures subject-specific information beyond the additional degrees of freedom provided by directionality alone. More generally, these results indicate that effective connectivity may provide a more informative representation of individual brain organization than standard FC, particularly in large-cohort settings and when relatively coarse parcellations are employed. This robustness is especially relevant for applications aimed at single-subject characterization, such as the prediction of cognitive traits or clinical phenotyping.

### EC-based network analysis

Having established the reliability and scalability of mOU-derived effective connectivity, we finally asked whether its directed nature can provide additional insight into the large-scale organization of brain networks. Unlike functional connectivity, EC allows us to characterize asymmetric interactions between regions and to investigate network properties that explicitly depend on directionality. We therefore examined the modular organization of the inferred EC, the distribution of positive and negative interactions across canonical resting-state networks, and the emergence of source–sink relationships and hierarchical organization across cortical regions. In Fig. 5a we show the group-level significant links of the averaged mou-EC matrix organized according to the 100-ROIs Schaefer parcellation. Regions are organized according to their assignment to one of the seven resting state networks (RSNs) in the Yeo 7-network partition, which aligns with a broad consensus about large-scale cortical networks [47], and includes the visual (Vis), somatomotor (SomMot), dorsal attention (DorsAttn), salience and ventral attention network (SalVentAttn), dorsoparietal control (Cont), and default mode network (Default), plus a set of orbitofrontal and temporal regions jointly classified as limbic (Limbic).

**Figure 5:**
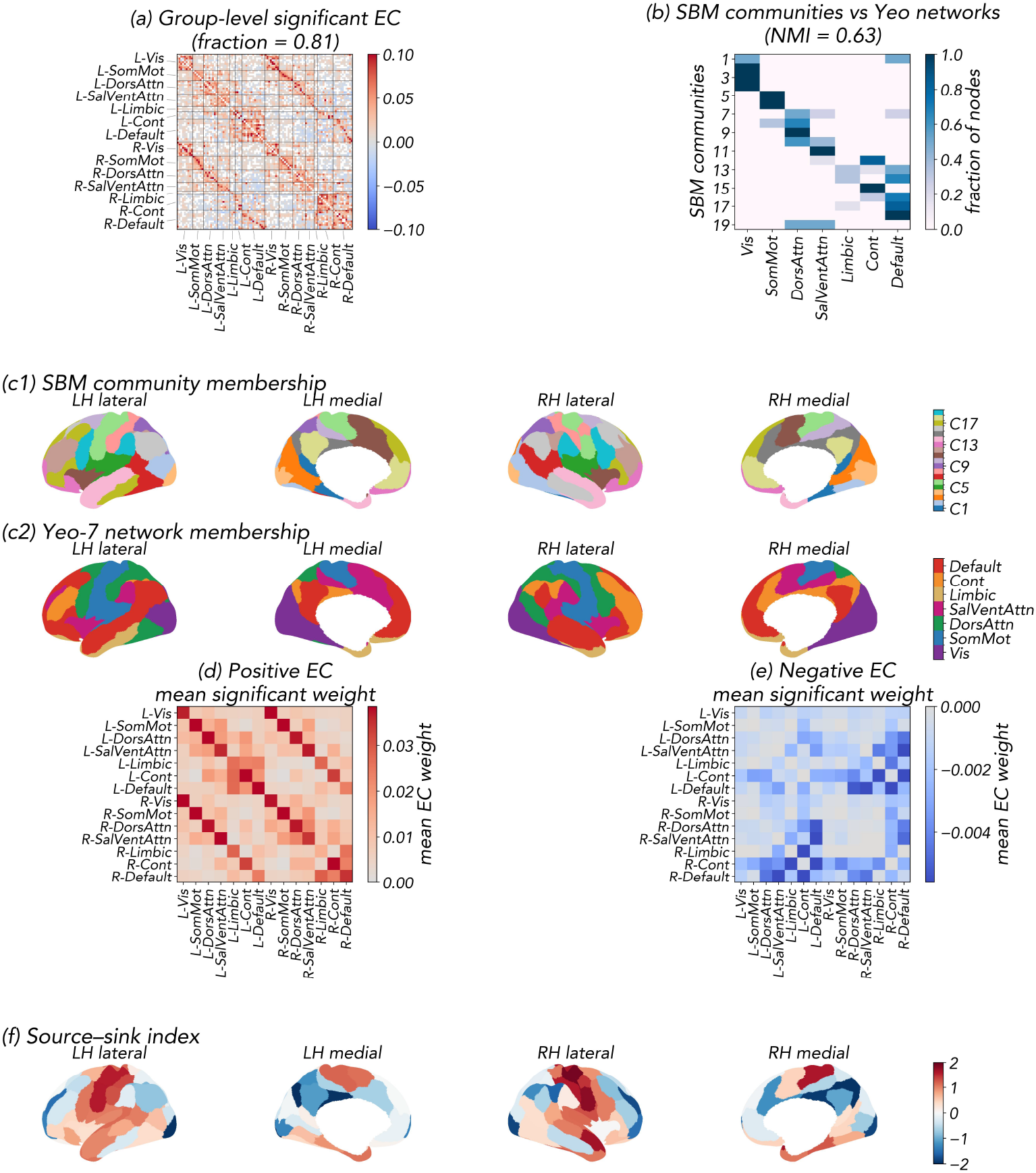
EC-based network analysis. **(a)** Group-level significant effective connectivity links. Directed connections were tested across subjects using one-sample statistical tests against zero and corrected for multiple comparisons. The matrix shows the group-averaged connectivity values restricted to statistically significant edges only, while the non-significant ones are highlighted in white. **(b)** Stochastic block model (SBM) community structure of the group-level EC in **(a)** compared to large-scale functional networks. Nodes are grouped into data-driven communities inferred from a weighted SBM fitted to the EC matrix. The resulting partition is compared against the Yeo resting-state networks, illustrating the correspondence between inferred and canonical network structure, which is highlighted by the Normalized Mutual Information (NMI) score. **(c1)** Stochastic block model (SBM) community membership visualized on the brain surface. **(c2)** Spatial distribution of the seven canonical Yeo functional networks across the Schaefer-100 cortical parcellation, shown as a reference for comparison with the independently inferred SBM communities in panel **(c1). (d)** Positive effective connectivity. Group-averaged directed connectivity restricted to positive weights, visualizing positive interactions between regions. **(e)** Negative effective connectivity. Group-averaged directed connectivity restricted to negative weights, reflecting negative interactions between regions. **(f)** Source–sink organization projected onto the cortical surface. Regional values of the source–sink index derived from the differential cross-covariance are projected onto a standard cortical template (Schaefer with 100 parcels), highlighting spatial gradients of network influence across the brain (positive values of the index indicate that a node is acting as a source, whereas negative values highlight sink roles).

We tested whether the presence of inter-network effective links would imply a network organization significantly different from the canonical one, the Yeo 7-network partition based on FC [48]. To this end, we applied a community detection algorithm (stochastic block modelling [49]) to cluster cortical regions based on effective connections. The algorithm found an optimal partition into 19 networks. Fig. 5b compares this partition with the canonical partition in 7 networks (merging their left- and right-hemisphere components). We found a strong overlap between the EC and FC-based networks, with a normalized mutual information of 0.63 between the two partitions. EC-based networks were all bilateral, and generally included both the left- and right-hemisphere components of the same FC-based networks. The similarity between the seven Yeo resting-state networks and the 19 communities identified by the stochastic block model (SBM) was quantified using normalized mutual information (NMI) [50]. Using the arithmetic-mean normalization, the resulting NMI was 0.629, while mininum- and maximum-entropy normalization gave NMI_min_ = 0.806 and NMI_max_ = 0.516. Overall, the results indicate substantial correspondence between the SBM communities and the canonical Yeo networks, despite the finer subdivision produced by the SBM (5c). The main difference between the two partitions is that some of the FC networks were split into separate sub-networks.

Then we studied the positive and negative maps of the EC. The mean significant value of positive and negative links within and between the main RSNs are shown in Fig. 5d,e. The highest value of significantly positive links is found within regions of the same networks, bilaterally (both intra- and inter-hemispherically), with the strongest connections being between homotopically symmetric regions. Significantly positive and negative inter-network links mirror known interactions between RSNs. Positive links between Vis and DorsAttn are consistent with the joint role of these networks in visuospatial control [51], and their connections through the posterior intraparietal sulcus and the middle temporal. Positive links between SomMot and SalVentAttn can be explained by the fact that the anterior cingulate cortex mediates interactions between these networks [52]. Coupling among attentional networks, the control network, and the default mode network reflects the presence of several connector hubs between these associative networks [53]. Positive Dorsattn-SalVentAttn connectivity may represent diffuse interactions of these two networks through several fronto-parietal connector hubs [54]. Positive links between Cont and Dorsattn/SalVentAttn reflect a coupling that is instrumental in attentional control [55]. Strong positive coupling between Default and Limbic is widely reported in the literature, which suggests that the boundaries between these two networks are not well defined [56]. The presence of negative links between Default and attentional networks (DorsAttn/SalVentAttn) is one of the most established features of resting state organization [4, 57]. Interestingly, Default and Cont exhibit both positive and negative connections, which is consistent with the simultaneous presence of cooperative and competitive interactions between these networks [58]. The main surprising feature emerging from the average EC is that inhibitory links are stronger inter-hemispherically than intra-hemispherically. This is at odds with the common belief that inter-hemispheric interactions are generally weaker than intra-hemispheric ones, as they must pass through the corpus callosum bottleneck [59] and evidence that between-network inhibition is primarily intra-hemispheric [60].

Finally, we exploited the differential cross-covariance *S* to characterize functional hierarchies in the cortex (Fig. 5f). For each node, we computed a group-level sink-source index (SSI), obtained by summing the column values of *S*, which isolates the net-direct flow contributions (see Methods). Positive SSI values indicate that a node predominantly acts as a *source* (driving other nodes), while negative SSI values pinpoint a *sink* (receiving input from other nodes). The group-level SSI score was positive for sensorimotor, ventrolateral prefrontal, orbitofrontal, secondary visual, and most temporal regions; it was negative for parietal, cingulate, medial prefrontal, primary visual, and lateral temporal regions. Overall, this asymmetry aligned with the transmodal-unimodal axis and the principal gradient of FC (correlation between SSI and principal gradient: −0.47, p-value *<* 10^*−*6^), and, moreover, the group-level SSI profile highly correlates with the one obtained from the sparse DCM EC matrices (see Fig S7, *r* = 0.725).

## Discussion

In this work, we systematically investigated the consistency, scalability, and subject-specific reliability of effective connectivity (EC) estimates obtained with the multivariate Ornstein–Uhlenbeck (mOU) model applied to resting-state fMRI data [34, 22, 14, 42, 41, 61, 62]. Using a large healthy cohort from the Human Connectome Project (*N* = 1013), we showed that mOU-based EC estimation provides a computationally efficient yet biologically plausible approximation of directed interactions at the whole-brain level. Importantly, our results demonstrate substantial agreement between mOU-derived EC and sparse Dynamic Causal Modeling (sparse DCM), despite the markedly different assumptions underlying the two frameworks.

A first contribution of this work is a technical improvement of the mOU-EC inference scheme. We showed that the standard procedure employed in previous implementations to estimate the global timescale parameter *τ*, based on the auto-correlation function of the signal, becomes unreliable when the latter is band-pass filtered in the conventional resting-state range (0.01–0.1 Hz). In this setting, short-lag autocorrelation estimates are strongly distorted by filtering effects, leading to excessively asymmetric and unstable solutions (Fig. 1). To address this issue, we examined a range of fixed timescales and selected a representative *τ* at the minimum of the group-average correlation between EC and the noise-weighted precision reference, within the range satisfying our convergence in at least 70% of participants (and initial-point exclusion criteria). This operational criterion identifies a reference within a convergent regime. The local minimum in correlation with the reversible reference potentially leaves greater scope for irreversible contributions within the convergent regime. The accompanying peak in correlation between antisymmetric EC and empirical lagged covariance suggests that these directional patterns are aligned with temporal asymmetries in the data (Fig. 1i).

Within this interval, EC estimates display a biologically plausible asymmetry and their connectivity patterns remain highly correlated across timescales (Fig. 1h, green). This result suggests that the mOU framework possesses a stable operating regime for filtered resting-state fMRI, without requiring fine-tuning to a single value of *τ*.

A second key result is the demonstration that mOU-based EC can be taken as a meaningful proxy for the more computationally expensive DCM-based EC. We compared mOU-EC estimates with those given by sparse DCM [30], one of the few currently available DCM variants capable of estimating whole-brain EC from resting-state fMRI while maintaining an explicit observation model for the hemodynamic response. Despite the fact that sparse DCM includes region-specific hemodynamic modeling and sparsity-inducing priors, while mOU relies on a substantially simpler linear stochastic formulation without an observation model, we observed high agreement between the two approaches both at the individual and at the group level (Fig. 2). Correlations between subject-level EC matrices were consistently high, and group-averaged EC estimates displayed remarkably similar large-scale organization. This finding indicates that in this healthy cohort the dominant large-scale directed interactions can already be captured by a simpler covariance-based dynamical model, without requiring the full complexity of nonlinear Bayesian inversion. Moreover, it suggests that, at least in healthy resting-state conditions, inter-regional variability in hemodynamic response function does not dramatically bias EC estimation at the temporal resolution of fMRI. This interpretation is supported by our analysis of HRF latencies, which revealed relatively limited regional variability across subjects (Fig. S2). While HRF variability did confound mOU-EC estimates, the bias remained relatively mild (Fig. S3). This finding partly diverges from the conclusion of Rangaprakash et al. [63], who argued in favor of a large confounding effect of HRF variability in FC estimation. This discrepancy could be due both to Rangaprakash et al.’s focus on FC rather than EC, and the specific method used for HRF inference (blind deconvolution) [64]. Future work may investigate possible inconsistencies between the two methods in HRF estimation more in depth.

A third main result of our analysis is that we provided evidence supporting the scalability of the mOU framework. While sparse DCM rapidly becomes computationally prohibitive beyond *∼* 90 regions, mOU inference can be efficiently extended to substantially larger parcellations. We demonstrated successful estimation with Schaefer atlases comprising up to 200 parcels, while preserving consistency across spatial scales (Fig 3). In particular, coarse-graining EC matrices inferred at high resolution yielded strong agreement with EC matrices estimated directly at lower resolution. This observation suggests that the inferred directed interactions are not merely the results of overfitting, but rather reflect stable mesoscale organizational features of brain dynamics [65, 66, 67]. The scalability of mOU is particularly relevant for future applications. Increasing spatial resolution is essential for studying fine-grained cortical hierarchies, local motifs of directed communication, and interactions between distributed functional systems [68, 69, 70]. Since computational cost grows rapidly in most DCM variants, scalable alternatives are required to bridge the gap between mechanistic modeling and high-resolution connectomics. Our results indicate that mOU provides a practical compromise between biological interpretability and computational tractability.

A fourth main finding of our study is the reliability of EC estimates at the individual subject level. Using repeated resting-state scans, we showed that mOU-derived EC matrices enable highly accurate “connectome fingerprints” [7], outperforming standard functional connectivity (FC), especially when large cohorts and relatively coarse parcellations are considered (Fig. 4). While FC is known to exhibit high within-subject stability, it also displays strong similarity across individuals, limiting discriminability as cohort size increases. In contrast, EC matrices appear to capture more individualized aspects of brain dynamics, likely because temporally lagged interactions encode additional dynamical constraints that are absent in non-lagged FC measures. The fact that EC fingerprinting remains highly accurate even with coarse parcellations suggests that effective connectivity contains subject-specific information at relatively large spatial scales. This robustness is particularly promising for translational applications such as clinical phenotyping, biomarker discovery, and individualized prediction [71]. Indeed, scalable and reliable EC estimation could enable the study of patient-specific alterations in directed communication patterns in neurological and psychiatric disorders [72, 73, 74, 75, 76].

Beyond consistency and scalability, the directed nature of EC enables analyses that are fundamentally inaccessible to FC. By exploiting asymmetry in the inferred connectivity matrices, we characterized source–sink organization across cortical regions and identified directed community structure through stochastic block modeling (Fig. 5). These analyses revealed structured patterns of directional influence that are compatible with known large-scale functional systems. Importantly, directed network measures such as in-strength, out-strength, and source–sink indices provide information about the hierarchy of information flow in the brain, potentially linking resting-state dynamics to theories of cortical processing hierarchies and predictive coding. Our results about flow asymmetries in the cortex are only partly in line with a previous analysis using Granger causality to identify asymmetries in the information flow between regions [77]: the main differences are that the primary visual and parietal cortices were identified as sources (rather than sinks). These differences may stem from technical differences between the two methods used to characterize asymmetries. While both methods are sensitive to 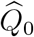 as well as *C* [78], the specific normalization choice used by [77] may have a strong impact on the results. A systematic comparison of the two methods is outside the scope of this article.

Several additional limitations should be acknowledged. First, the mOU model is a linear stochastic approximation and cannot capture nonlinear or state-dependent neuronal interactions. It assumes a common regional decay timescale and does not explicitly model region-specific hemodynamics. Second, the analysis was restricted to cortical parcels, a single parcellation family, one phase-encoding direction per session, and one preprocessing strategy. Moreover, the comparison with sparse DCM involved a subset of participants and off-diagonal connectivity only, because the two models treat diagonal terms (i.e., self-decay time constants) differently. Third, neither cross-model agreement nor cross-session reproducibility establishes biological ground truth. Validation against electrophysiological [79] or perturbational measurements [80] will be essential for determining the neuronal specificity of the inferred directional interactions. Finally, the proposed effective connectivity framework was developed and validated for healthy populations. Its application to neurological or psychiatric disorders affecting cerebral hemodynamics should therefore be undertaken with caution and only after verifying that the underlying assumptions remain valid for the specific pathology under investigation.

Future work could investigate more principled approaches to fix timescales in mOU, including region-specific timescale estimation strategies, and integrate structural connectivity constraints in EC estimation [81, 82], potentially improving the biological realism of the model. Furthermore, it could focus on neural pathologies or task-evoked dynamics, which may require region-specific hemodynamic modeling and/or richer models incorporating external inputs [83] and context-dependent modulation. Finally, combining scalable EC estimation with machine learning techniques may allow developing individualized biomarkers based on directed brain dynamics [84].

In conclusion, mOU provides a computationally efficient framework for estimating whole-brain directed coupling from resting-state fMRI in healthy adults. For filtered data, we identify a convergent range of decay timescales and select the solution as the global minimum of the reversible reference, while also assessing that the connectivity patterns across that range are robust. Within this regime, reduced alignment with the reversible reference, together with stronger correspondence to empirical lagged asymmetry, suggests room for irreversible contributions that reflect directional structure in the data. Under the conditions studied here, its inter-regional estimates show substantial correspondence with sparse DCM, remain stable across spatial scales, and capture reproducible individual and network-level features. These properties make mOU a promising tool for large-scale investigations of directed brain organization, provided that its hemodynamic assumptions and model dependence are explicitly considered when extending the framework to new populations and experimental settings.

## Methods

### The mOU model

#### Dynamical model

The multivariate Ornstein-Uhlenbeck (mOU) model proposed by Gilson and colleagues provides a linear stochastic framework for describing whole-brain dynamics in terms of effective connectivity and noise propagation. The temporal evolution of the regional activity vector **x**(*t*) is governed by the stochastic differential equation:

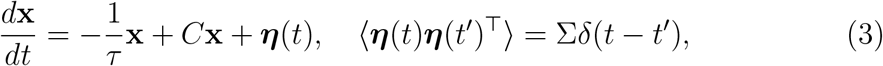

where *C* is the effective connectivity (EC) matrix, 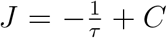 is the Jacobian of the dynamics, Σ is the noise covariance, and ***η***(*t*) is Gaussian white noise. The parameter *τ* acts as a time scale for the system, and it globally impacts the dynamics.

Under stationarity, the covariance matrix at zero time-lag, denoted *Q*_0_, satisfies the continuous-time Lyapunov equation:

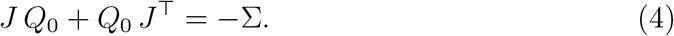

Solving Eq. 4 provides the model-predicted zero-lag covariance, while the lagged covariance at time *t*_lag_ is given by:

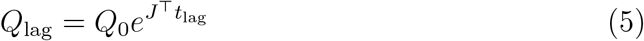

Note that in principle, from the formula above, one could directly obtain an estimate 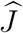 through the empirical covariance and lagged covariance 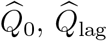.

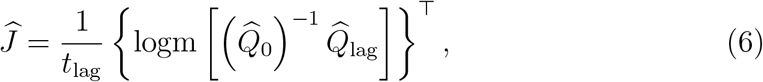

where logm is the matrix logarithm. Yet, instabilities in matrix inversion and matrix logarithm extraction make this method very sensitive to the noise in 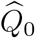 and 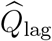, leading to the appearance of non-real entries in 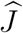, hence in *C*. For this reason, Gilson et al. [34] developed an optimization method to overcome this issue.

#### Estimation of effective connectivity via Lyapunov optimization

We denote by 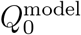 and 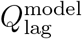 respectively the covariance matrices of the model at zero time-lag and at a time-lag *t*_lag_. Given empirical covariances 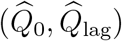, we define the errors

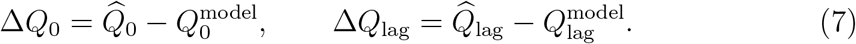

We want to update *C* to obtain the desired changes for the model covariances *Q*_0_ and *Q*_lag_. Moreover, we z-scored the time series, in accordance with rescaling practices that are recommended when dealing with DCM models [85]. Hence, in practice, the covariance matrices we consider are in fact correlation matrices. In the update formulas below, *Q*_0_ and *Q*_lag_ denote the current model covariances.

The Lyapunov optimization (LO) update for the Jacobian **J** is derived from the gradient of the loss

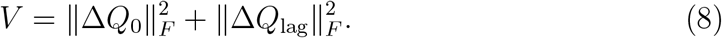

In particular, letting

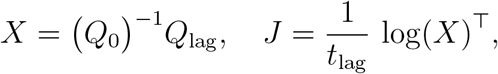

we perform an implicit differentiation with respect to *J* and *X*, yielding the corresponding update of the Jacobian:

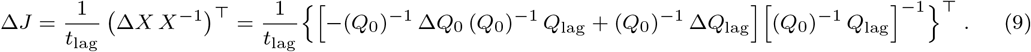

Then, using the identity *Q*_lag_ = *Q*_0_ exp(*J* ^*⊤*^*t*_lag_), one can also write:

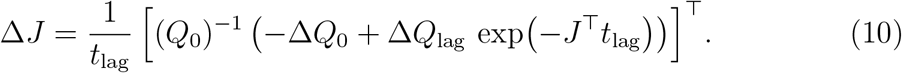

The connectivity update will thus be: Δ*C*_*ij*_ = *η*_*C*_Δ*J*_*ij*_, for *i* ≠ *j*., i.e., a gradient descent to minimize the loss in eq. 8. The diagonal elements of *J* are not included in this optimization loop, but are fixed beforehand. We have 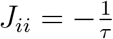. In [34, 41], *τ* is simply estimated by fitting an exponential only to the first two, or three, points of the autocorrelation function.

Moreover, for the 200-region parcellation, for which the range of parameters yielding stable convergence was narrower, we additionally employed the optional *L*_2_ regularization implemented in the mOU optimization procedure. Following the connectivity update introduced by Gilson et al., the regularized update can be written as

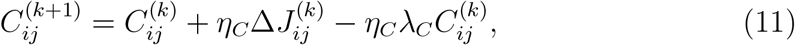

where *η*_*C*_ is the learning rate and *λ*_*C*_ corresponds to the *regul*_*C*_ parameter. The additional term provides *L*_2_ shrinkage of the EC weights and was used for the 200-region parcellation to improve the numerical stability of the optimization and facilitate convergence. For the lower-resolution parcellations, *λ*_*C*_ was set to zero. Eq. 10 contains a free parameter, *t*_lag_. While in principle any choice of *t*_lag_ would be correct, in [34], Gilson and colleagues show that that the method is, in practice, most efficient for a specific choice of *t*_lag_. By extensively studying the relationship between *t*_lag_ and *τ* through numerical simulations, they noted that *t*_lag_ has to be chosen to approximately match *τ* . In fact, for *t*_lag_ *≪ τ* the EC matrices obtained are highly symmetric, and directionality is lost; on the other hand, for *t*_lag_ *≫ τ* the matrices are highly asymmetric, but also random and noisy (high errors in recovering *C* and *Q*_0_ / *Q*_lag_). In fMRI studies, one generally sets *t*_lag_ = 1 TR [14, 41]).

Finally, to take properly the effect of Σ, in [14, 41] the Σ update has been adjusted from the heuristic update in the first study [34] to a gradient descent:

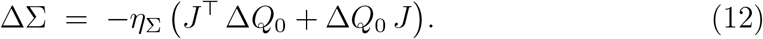

#### Computational cos

mOU inversion took an average time of 0.76 seconds per subject to estimate an EC with 100 parcels; this computing time increases just to 15.8s if one also counts the grid search across 20 different values of *τ*).

### Sparse DCM

In this work we use the sparse DCM approach by Prando et al.[30]. This DCM variant implements a sparsity-inducing mechanism that automatically prunes irrelevant connections, thereby avoiding the need to perform a selection between competing network structures.

The model includes two layers: (i) a coupled ODE system modeling neuronal activation **x**(*t*), and (ii) a mapping from neuronal activity **x**(*t*) to the BOLD fMRI signal **y**(*t*) (hemodynamic response):

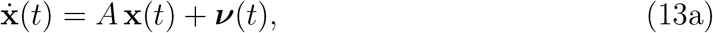

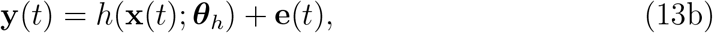

where **x**(*t*) = [*x*_1_(*t*) … *x*_*n*_(*t*)]^*T*^ is the hidden neuronal activity across *n* regions, *A* is the effective connectivity matrix (with entry *A*_*ij*_ representing the influence from *j* to *i*), ***v***(*t*) is a stochastic driving term, *h*(· ; ***θ***_*h*_) is the hemodynamic transformation parameterized by ***θ***_*h*_, and **e**(*t*) *∼ N* (0, *R*) is Gaussian observation noise with covariance *R*. Note that the matrix *A* in sparse DCM corresponds to the Jacobian matrix *J* in mOU. Contrary to mOU, sparse DCM also estimates self-connections *A*_*ii*_, which are always negative, thus corresponding to (region-specific) self-decay terms,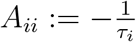.

To simplify the estimation, two main steps are taken. First, due to the low temporal resolution of fMRI (sampling time *T*_*R*_ ≈ 0.5–3s), the continuous dynamics are discretized:

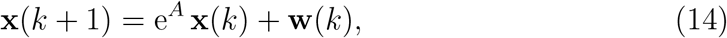

where **w**(*k*) is white Gaussian noise with covariance *Q*. Second, the hemodynamic response is linearized using a finite impulse response (FIR) model:

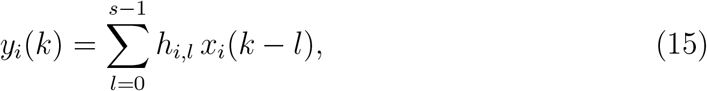

with FIR coefficients *h*_*i*_ = [*h*_*i*,0_, …, *h*_*i,s−*1_]^*T*^.

Combining these, the model becomes linear stochastic:

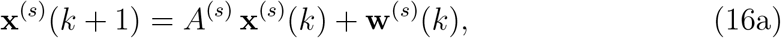

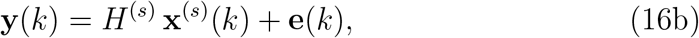

where **x**^(*s*)^(*k*) = [**x**^*T*^ (*k*) … **x**^*T*^ (*k*− *s* + 1)]^*T*^, *A*^(*s*)^ and *H*^(*s*)^ contain the EC and FIR parameters, and **w**^(*s*)^ has covariance *Q*^(*s*)^. All parameters *θ* = {*A, H, Q, R*} are estimated in a Bayesian framework with factorized priors:

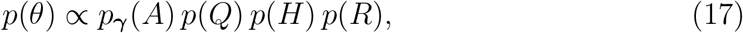

where *p*(*Q*) and *p*(*R*) are uninformative, *p*(*H*) is Gaussian with fixed mean and variance, and *p*_***γ***_(*A*) is a sparsity-inducing prior

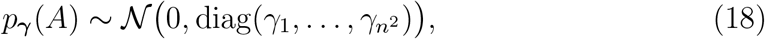

with hyperparameters *γ*_*i*_ estimated by marginal likelihood maximization to induce sparsity in *A*. Parameters are obtained via a maximum a-posteriori estimate using an expectation–maximization algorithm. The resulting effective connectivity matrix is thereby sparse, facilitating more interpretable whole-brain EC estimation from resting-state fMRI.

#### Computational cost

Even when using a parcellation with lower resolution, and although the model was run on a high-performance computing cluster, sparse DCM estimation typically required at least 6 hours of computing time per subject (vastly outperformed by mOU in terms of computational efficiency).

### Datasets

We used 1013 subjects from the Human Connectome Project (HCP) 1200 subjects release [35]. We used the left-right (LR) runs from the first session (REST 1) of resting state fMRI data, and from the second session (REST 2) to perform fingerprinting. Time series had 1200 time points with a TR of 0.72 sec, meaning ≈ 15 mins duration. The full description of the imaging parameters and minimal preprocessing pipeline is given in Ref. [86]. In short, after correction for motion, gradient, and susceptibility distortions the fMRI data was aligned to an anatomical image. The aligned functional image was corrected for intensity bias, demeaned, and projected to a common surface space, which resulted in a cifti-file. Artifacts were removed though Independent Component Analysis (ICA) using the FSL’s MELODIC tool paired with the FMRIB’S ICA-based X-noisefilter. No additional global signal regression was applied. All fMRI data were filtered between 0.01 and 0.1 Hz to retain the relevant frequency range for further analyses of the BOLD signal. Functional data can be mapped to different spatial resolutions using the Schaefer parcellation [87], which optimizes local gradient and global similarity measures of the fMRI signals. Regional time series were extracted using Workbench Command provided by the HCP.

### Parcellations

#### From 100-parcels to 62-parcels resolution

The cortical parcellation employed in this study comprises 62 regions derived from the Schaefer 100-area atlas organized into 7 functional networks. The decision to cluster the original atlas was performed in previous works [40, 39] and stems from the necessity to maintain the computational load of sparse DCM at a manageable level. The clustered atlas was created by merging areas that were already functionally grouped in the atlas, as detailed by [39], while keeping other areas separate. Specifically, a consensus clustering procedure was employed to reduce the number of cortical parcels derived from the Schaefer atlas. This framework employs both base and consensus clustering methods to achieve robust and stable clusters. Initially, various groupings are generated for each individual dataset using a base clustering method. Then, consensus clustering is applied to these groupings to produce stable clusters across individuals. To account for hemodynamic differences across spatially distant parcels, this procedure is selectively performed on subsets of adjacent cortical regions within the same functional network. This additional constraint ensures that only functionally homogeneous and spatially contiguous parcels are grouped together. For sets composed of only two contiguous ROIs, the objective criteria for determining the optimal number of clusters cannot be applied. In these cases, grouping is performed if the pair Pearson’s correlation is, on average, greater than the mean correlation within other clusters. This approach ensures the hemodynamic consistency of each cluster, reflecting the whole-brain vascular complexity.

#### From 200-parcels to 100-parcels resolution

To establish a correspondence between the Schaefer 200-parcel and 100-parcel atlases, we derived a vertex-wise mapping based on shared cortical vertices. Both atlases are defined in the same surface space, allowing a direct comparison at the vertex level.

The mapping from 200 to 100 parcels was defined using a majority-vote criterion: each 200-parcel region was assigned to the 100-parcel region that occurred most frequently among its shared vertices. This procedure ensured that the mapping was driven exclusively by vertices that were unambiguously present in both parcellations and minimized the influence of unmatched or boundary vertices.

This vertex-wise majority assignment resulted in a deterministic many-to-one mapping from the 200-parcel atlas to the 100-parcel atlas, which was subsequently used to aggregate region-level measures across spatial scales.

### Differential cross-covariance and source-sink index

Considering, as before, the multivariate Ornstein–Uhlenbeck process described

by

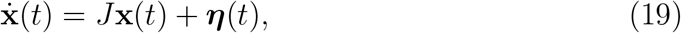

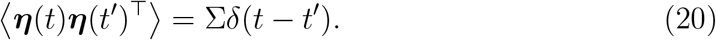

At stationarity, the zero-lag covariance matrix *Q*_0_ satisfies the continuous Lyapunov equation

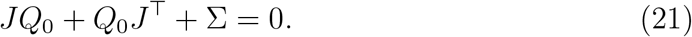

Note that the Python package pyMOU, implementing the mOU framework [34, 41], adopts a row-vector convention, 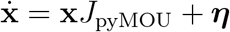 . Accordingly, its stationary covariance satisfies

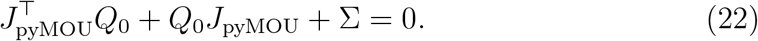

Here we use the column-vector convention, with 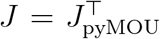, so that the same equation reads 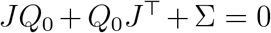. We therefore transpose the Jacobian returned by pyMOU before computing the differential cross-covariance below.

The differential covariance matrix quantifies the covariance between the temporal derivative of one variable and the instantaneous state of another variable [88, 89], and is defined as [38]

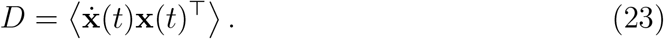

For linear dynamics, it is related to the effective connectivity and the stationary covariance by

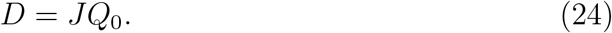

The differential cross-covariance can be decomposed into symmetric and anti-symmetric components:

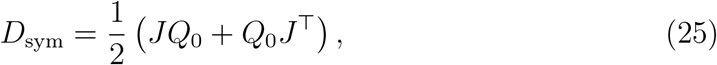

and

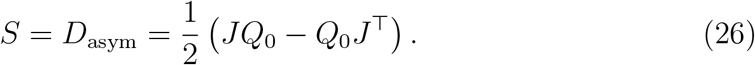

From the Lyapunov equation, the symmetric component is

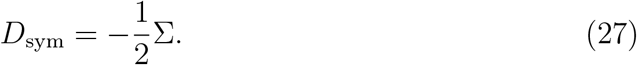

The antisymmetric component is called differential cross-covariance [38] (we will refer to it as *S*) and contains the part of the dynamics that is not directly determined by the noise covariance. Detailed balance holds when

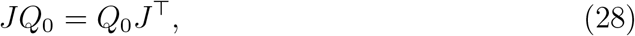

which implies

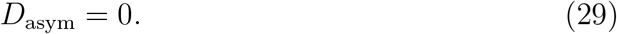

In this condition, the stationary dynamics is time-reversible and the stationary probability current vanishes. Conversely, a non-zero antisymmetric differential cross-covariance *S* indicates broken detailed balance and irreversible dynamics in a non-equilibrium steady state. It is therefore associated with circulating probability currents in the state space, even though the probability distribution remains stationary. From 24 it can be seen that J can be expressed as a sum of two contributions:

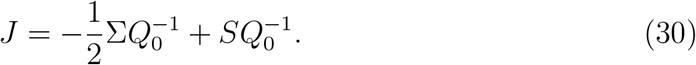

The differential cross-covariance isolates the time-irreversible component of the dynamics, namely the pairwise directional imbalance that cannot be explained by the symmetric, detailed-balance contribution. Therefore, the column sum

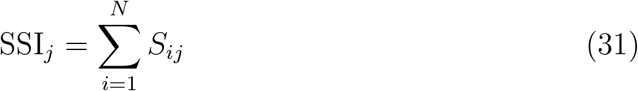

provides a signed source–sink index, measuring the net imbalance between dynamical flow leaving and entering node (j). A positive value indicates a prevalence of outgoing flow, identifying the node as a source, whereas a negative value indicates a prevalence of incoming flow, identifying it as a sink, according to the adopted matrix convention.

### HRF latencies calculation

Hemodynamic response functions (HRFs) estimated from sparse DCM were used to extract peak-related temporal features for each subject and signal. For each signal, the HRF time course was interpolated over a dense temporal grid of 1000 points spanning the original sampled time window (16 seconds). A quadratic interpolation was used to obtain a smoother estimate of the response profile.

The HRF amplitude was defined as the maximum value of the interpolated HRF, and the HRF latency was defined as the time point at which this maximum occurred. To estimate the full width at half maximum (FWHM), the half-maximum threshold was computed as half of the peak amplitude. The two threshold crossings surrounding the peak were then identified, one before and one after the peak. FWHM was calculated as the temporal distance between the right and left half-maximum crossings.

### Stochastic Block model communities

We inferred a stochastic block model (SBM) [49] to identify communities in the network described by the weighted and directed effective connectivity. Indeed, SBM is known to overcome the known problems of other community detection algorithms, which maximize modularity, since the latter will end up finding communities even if they are not truly present in the network [90]. Moreover, this formulation naturally supports *directed weighted networks*, in contrast to many community detection approaches that assume symmetry (e.g., spectral clustering or modularity maximization on undirected graphs). The stochastic block model (SBM) was inferred through Markov Chain Monte Carlo (MCMC) sampling. A directed (or undirected) weighted graph *G* = (*V, E*) was constructed from *EC*, where edges exist for *EC*_*ij*_ *≠*0 and are assigned weights:

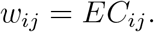

The SBM assumes a block partition *b* and block-specific connectivity parameters *θ*, and is fitted by minimizing the description length:

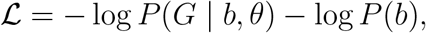

which provides a principled Bayesian model selection criterion that automatically balances model complexity and goodness of fit. An initial partition was obtained by minimizing the SBM description length. Posterior exploration was performed via MCMC sampling [91] for *T* iterations, with thinning parameter *τ* (i.e., `iter_skip`). At each iteration, we stored the partition *b*^(*t*)^ and the corresponding model entropy

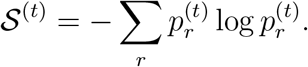

This yields a posterior ensemble of partitions:

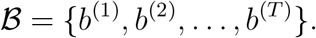

A consensus partition *b*^*\**^ was obtained via mode clustering over the posterior samples:

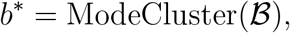

providing a robust estimate of community structure that accounts for degeneracy in the SBM posterior. The method returns the consensus partition *b*^*\**^, the block-to-block connectivity matrix *e*_*rs*_, and the entropy trace {S^(*t*)^}.

#### Normalized mutual information

Let *Y* and *S* denote two partitions, respectively. NMI can be computed as

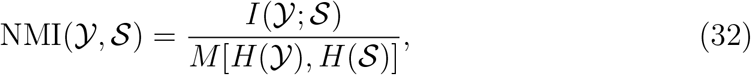

where *I*(*Y*; *S*) is their mutual information, *H*(*Y*) and *H*(*S*) are the entropies of the two partitions, and *M* denotes the normalization function. The default choice is arithmetic-mean normalization, 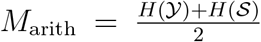. Minimum- and maximum-entropy normalizations (obtained using *M* = min {*H*(*Y*, *H*(*S*)} and *M* = max {*H*(*Y*, *H*(*S*)} are also commonly used. The larger value obtained with the minimum normalization reflects the use of the lower-entropy partition as the reference scale and therefore quantifies how much of the information contained in the simpler partition is captured by the other partition. Conversely, the maximum normalization uses the higher-entropy partition as the denominator and is therefore more conservative when the two partitions differ substantially in resolution.

## Acknowledgments

This project has been supported by #NEXTGENERATIONEU (NGEU) and funded by the Ministry of University and Research (MUR), National Recovery and Resilience Plan (NRRP), project MNESYS (PE0000006) - A Multiscale integrated approach to the study of the nervous system in health and disease (DN. 1553 11.10.2022).

## Data and code availability

The neuroimaging data analyzed in this study are available through the Human Connectome Project (HCP; https://www.humanconnectome.org/study/hcp-young-adult). The code used for the calculation of the mOU EC and the generation of the figures is publicly available at https://github.com/benedetta-mariani/mOU-EC-estimation. The underlying pyMOU package is available at https://github.com/mb-BCA/pyMOU.

## Supplementary Information

**Figure S1:**
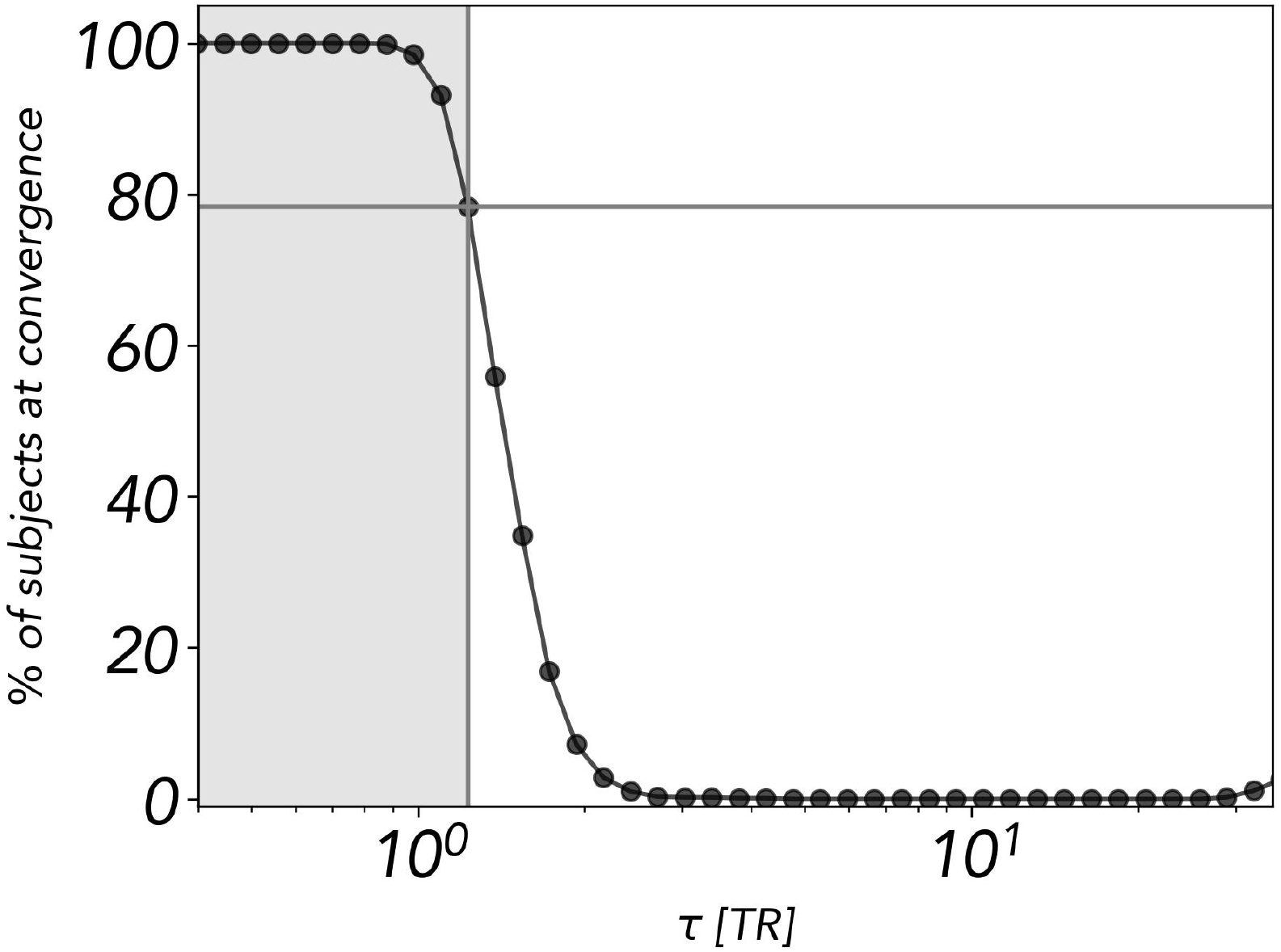
Percentage of subjects for which the mOU optimization converged as a function of the timescale parameter *τ* . For each value of *τ*, the plot shows the percentage of subjects for which the EC estimation reached a stable solution according to the predefined convergence criterion. Convergence progressively decreased for larger values of *τ* ; at *τ* = 1.23 TR, 78% of subjects still reached convergence.

**Figure S2:**
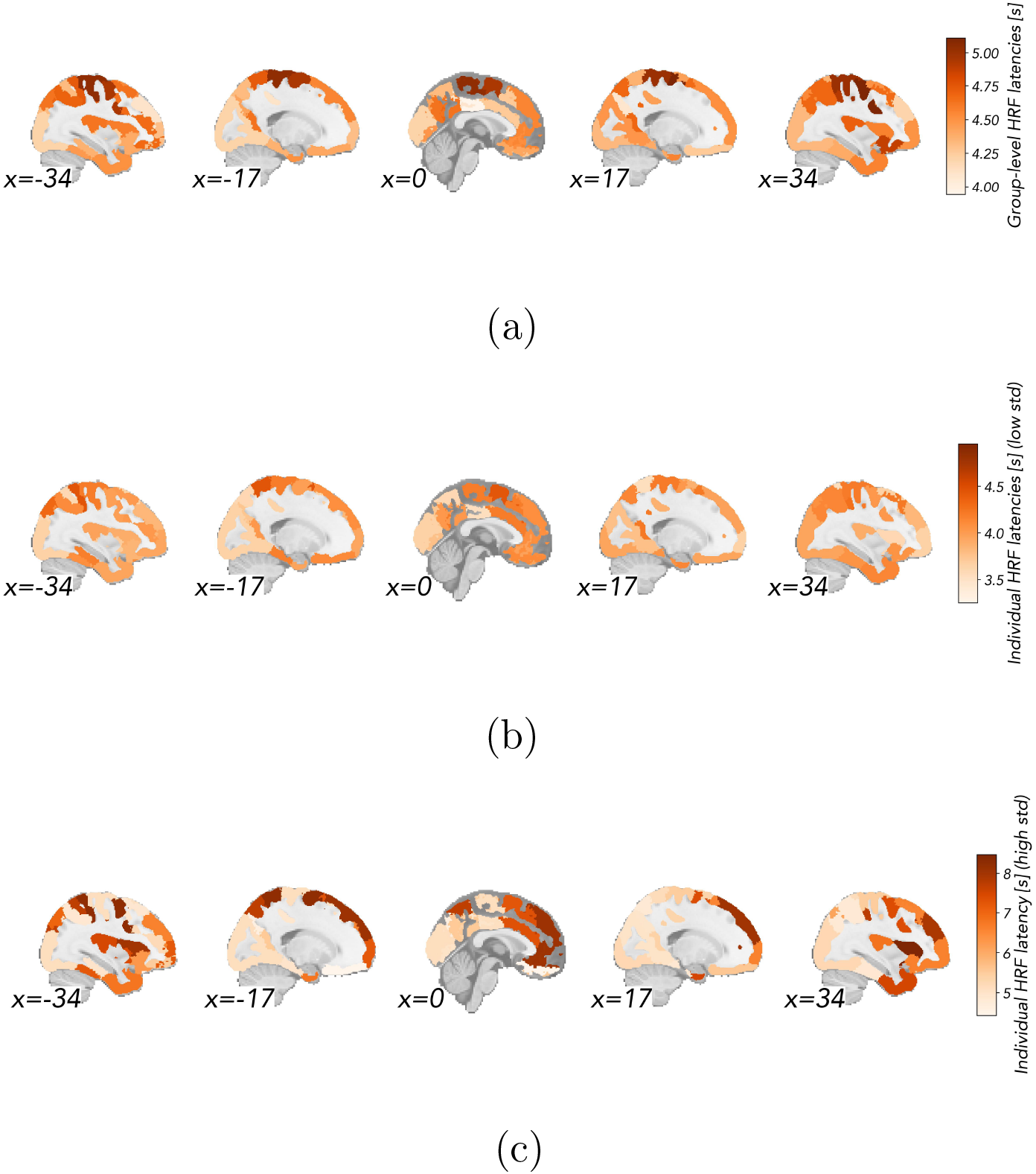
(a) Group level HRF latencies. We found only moderate differences in HRF latencies across regions - typically, in the range [3.6 s, 6.4 s] (5th and 95th percentile of the latencies distribution, including all the subjects and all the regions) **(b) Single-subject HRF latencies**, referring to the subject with the lowest standard deviation of the latencies across brain regions. **(c) Single-subject HRF latencies**, referring to the subject with the highest standard deviation of the latencies across brain regions.

**Figure S3:**
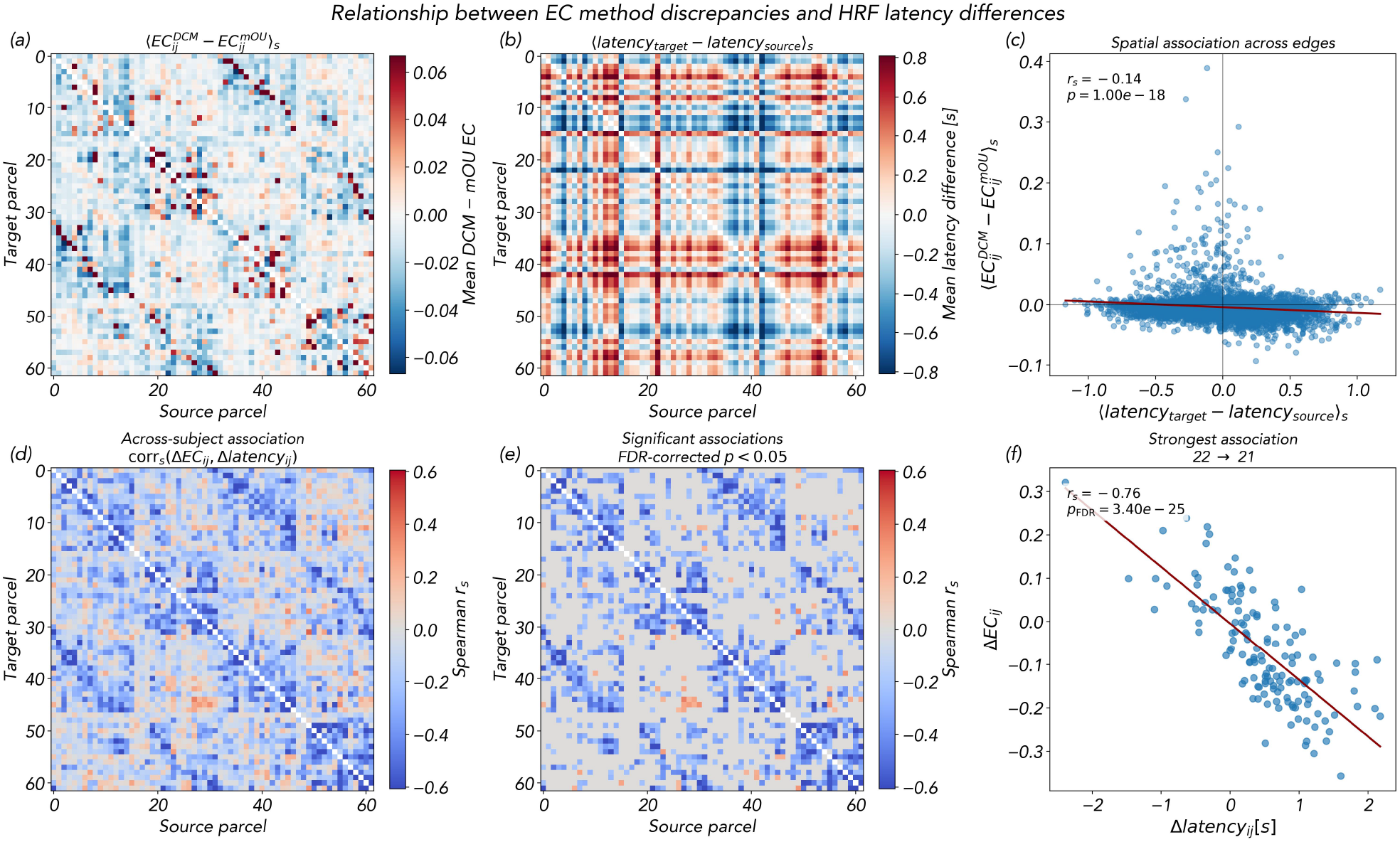
Relationship between regional HRF latency differences and discrepancies between mOU- and sparse DCM-based effective connectivity estimates. **(a)** Group-average signed difference between sparse DCM and mOU effective connectivity estimates, computed for each directed edge as 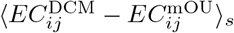. Positive values indicate larger EC estimates for sparse DCM, whereas negative values indicate larger estimates for mOU. **(b)** Group-average signed inter-regional HRF latency difference, *⟨L*_target_ − *L*_source_*⟩* _*s*_. Positive values indicate that the target region exhibits a later HRF peak than the source region. **(c)** Spatial association across directed edges between the mean signed HRF latency difference and the mean signed EC discrepancy. Each point represents one directed edge, the solid line indicates a linear fit for visualization, and the Spearman correlation coefficient is reported in the panel. **(d)** Edge-wise Spearman correlation, computed across subjects, between the subject-specific EC discrepancy and the corresponding subject-specific HRF latency difference. **(e)** Directed edges for which the across-subject association remained significant after false-discovery-rate correction. **(f)** Subject-level relationship for the edge showing the strongest absolute Spearman correlation. Each point represents one subject; the solid line indicates a linear fit for visualization, and the Spearman correlation coefficient and FDR-adjusted *p*-value are reported in the panel. Matrix rows and columns represent target and source regions, respectively. We systematically find a negative correlation between Δ*C*_*ij*_ and Δ*ℓ*_*ij*_ across subjects, in most of the EC links for which we observe a significant effect.

**Figure S4:**
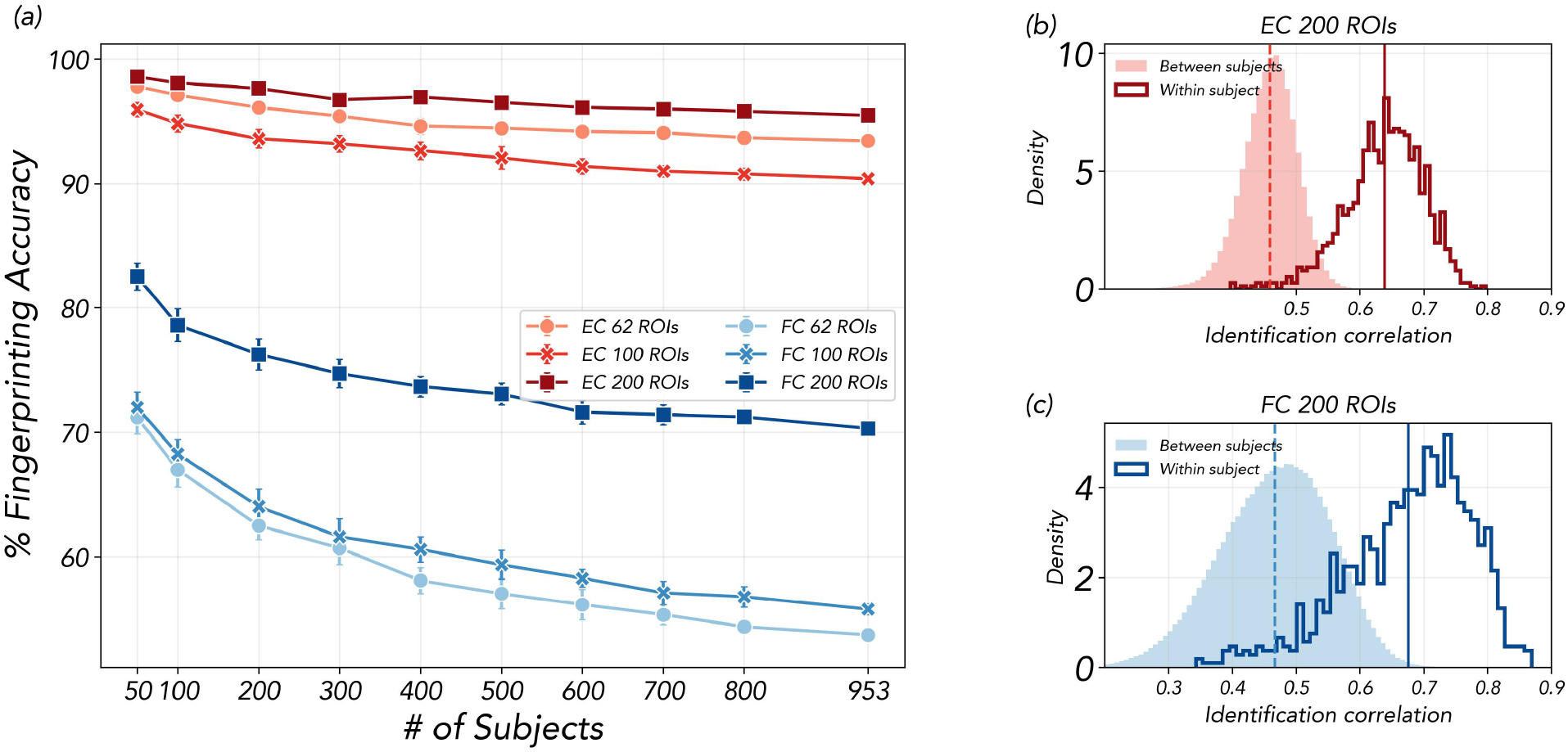
Fingerprinting performance of symmetrized effective connectivity and functional connectivity across parcellation scales. **(a)** Identification accuracy between REST1 and REST2 as a function of the number of subjects for symmetrized effective connectivity (EC) and functional connectivity (FC), evaluated using parcellations with 62, 100, and 200 regions of interest. Error bars indicate variability across repeated subsampling realizations. **(b–c)** Distributions of within-subject correlations, corresponding to the diagonal elements of the identification matrix, and between-subject correlations, corresponding to its off-diagonal elements, for symmetrized EC and FC, respectively. Symmetrizing EC matches its number of independent off-diagonal elements to that of FC, showing that the higher fingerprinting performance of EC cannot be explained solely by its larger number of directed parameters.

**Figure S5:**
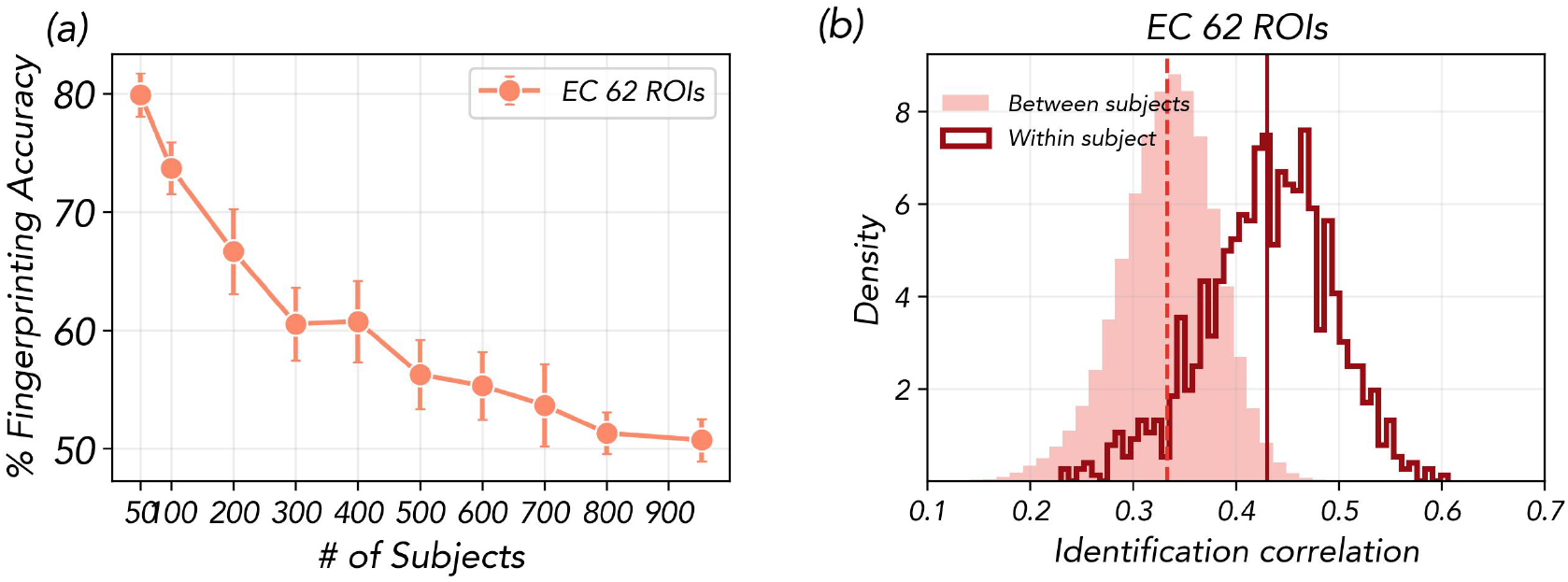
Effective-connectivity fingerprinting using unfiltered resting-state fMRI data. Effective connectivity (EC) was estimated separately from REST1 and REST2 using unfiltered fMRI time series and the 62-region parcellation. **(a)** Subject-identification accuracy as a function of cohort size. Points show the mean accuracy across randomly selected subject subsets, and error bars indicate three times the standard error of the mean. **(b)** Distributions of the diagonal (within-subject) and off-diagonal (between-subject) entries of the REST1–REST2 identifiability matrix. Vertical lines indicate the corresponding distribution means. Compared with EC estimated from temporally filtered data, EC estimated from unfiltered data yielded lower fingerprinting accuracy and lower self-identifiability, indicating reduced consistency of individual EC patterns across independent resting-state runs.

**Figure S6:**
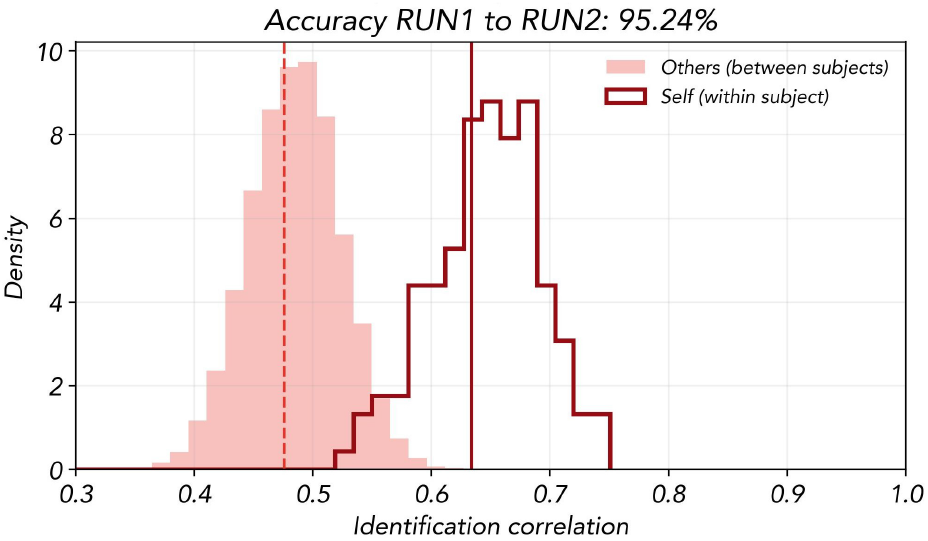
Subject fingerprinting using DCM connectivity. Within-subject (self) and between-subject (others) identification correlation distributions across two runs. Pearson correlations were computed from directed off-diagonal connections among 62 ROIs for the 144 subjects available in both runs. Solid and dashed vertical lines indicate the mean within-subject and between-subject correlations, respectively. Each subject in RUN1 was matched to the subject with the highest correlation in RUN2, yielding an identification accuracy of 95.24%.

**Figure S7:**
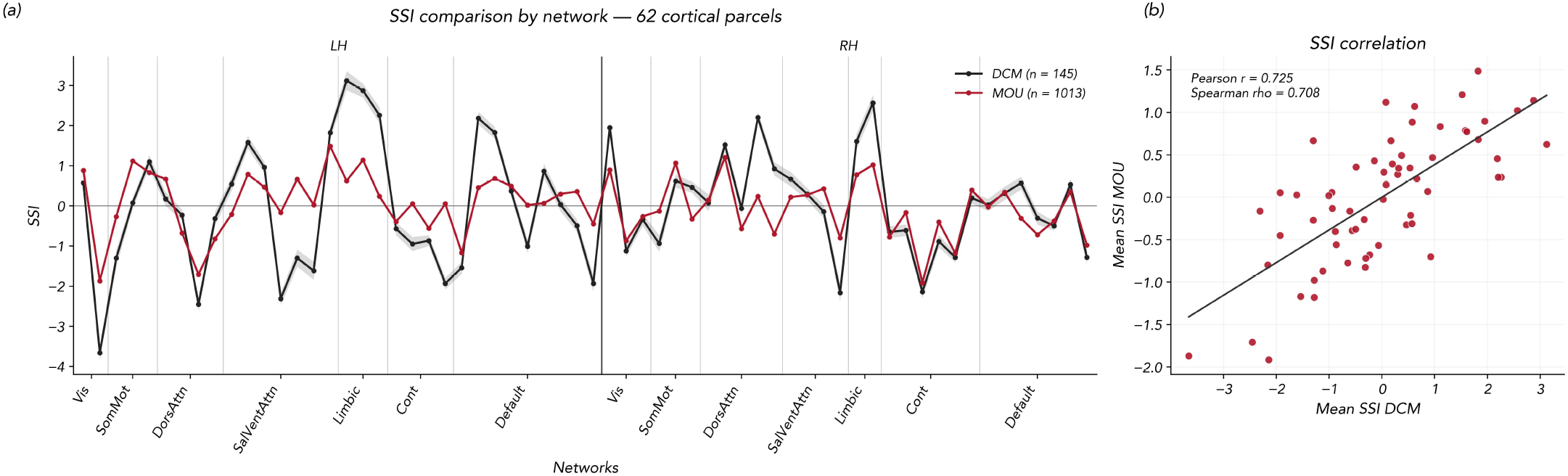
Group-level comparison of cortical SSI between DCM and mOU. **(a)** Mean source–sink index (SSI) across 62 cortical parcels, grouped by functional network, for DCM and mOU. Shaded bands indicate the standard error of the mean across subjects. **(b)** Correlation between the group-averaged regional SSI profiles from the two models. Each point represents one cortical parcel (Pearson’s *r* = 0.725; Spearman’s *ρ* = 0.708).

## Notes

### Competing Interest Statement

The authors have declared no competing interest.

## References

[1] Olaf Sporns, Dante R Chialvo, Marcus Kaiser, and Claus C Hilgetag. Organization, development and function of complex brain networks. Trends in cognitive sciences, 8(9):418–425, 2004.

[2] Danielle S Bassett and Edward T Bullmore. Human brain networks in health and disease. Current opinion in neurology, 22(4):340, 2009.

[3] Steven L Bressler and Vinod Menon. Large-scale brain networks in cognition: emerging methods and principles. Trends in cognitive sciences, 14(6):277–290, 2010.

[4] Michael D Fox, Abraham Z Snyder, Justin L Vincent, Maurizio Corbetta, David C Van Essen, and Marcus E Raichle. The human brain is intrinsically organized into dynamic, anticorrelated functional networks. Proceedings of the National Academy of Sciences, 102(27):9673–9678, 2005.

[5] Michael D Fox and Michael Greicius. Clinical applications of resting state functional connectivity. Frontiers in systems neuroscience, page 19, 2010.

[6] Jonathan D Power, Alexander L Cohen, Steven M Nelson, Gagan S Wig, Kelly Anne Barnes, Jessica A Church, Alecia C Vogel, Timothy O Laumann, Fran M Miezin, Bradley L Schlaggar, et al. Functional network organization of the human brain. Neuron, 72(4):665–678, 2011.

[7] Emily S Finn, Xilin Shen, Dustin Scheinost, Monica D Rosenberg, Jessica Huang, Marvin M Chun, Xenophon Papademetris, and R Todd Constable. Functional connectome fingerprinting: identifying individuals using patterns of brain connectivity. Nature neuroscience, 18(11):1664–1671, 2015.

[8] Karl J Friston. Functional and effective connectivity: a review. Brain Connectivity, 1(1):13–36, 2011.

[9] Karl J Friston, Lee Harrison, and Will Penny. Dynamic causal modelling. Neuroimage, 19(4):1273–1302, 2003.

[10] Edmund T Rolls, Gustavo Deco, Chu-Chung Huang, and Jianfeng Feng. The human language effective connectome. NeuroImage, 258:119352, 2022.

[11] Derek Evan Nee and Mark D’Esposito. The hierarchical organization of the lateral prefrontal cortex. Elife, 5:e12112, 2016.

[12] Weidong Cai, Srikanth Ryali, Ramkrishna Pasumarthy, Viswanath Talasila, and Vinod Menon. Dynamic causal brain circuits during working memory and their functional controllability. Nature Communications, 12(1):3314, 2021.

[13] Kyesam Jung, Karl J Friston, Chongwon Pae, Hanseul H Choi, Sungho Tak, Yoon Kyoung Choi, Bumhee Park, Chan-A Park, Chaejoon Cheong, and Hae-Jeong Park. Effective connectivity during working memory and resting states: A dcm study. NeuroImage, 169:485–495, 2018.

[14] Matthieu Gilson, Gustavo Deco, Karl J Friston, Patric Hagmann, Dante Mantini, Viviana Betti, Gian Luca Romani, and Maurizio Corbetta. Effective connectivity inferred from fmri transition dynamics during movie viewing points to a balanced reconfiguration of cortical interactions. Neuroimage, 180:534–546, 2018.

[15] Seonghoon Kim, Hyun Seok Moon, Thanh Tan Vo, Chang-Ho Kim, Geun Ho Im, Sungho Lee, Myunghwan Choi, and Seong-Gi Kim. Whole-brain mapping of effective connectivity by fmri with cortex-wide patterned optogenetics. Neuron, 111(11):1732–1747, 2023.

[16] Maksim G Sharaev, Viktoria V Zavyalova, Vadim L Ushakov, Sergey I Kartashov, and Boris M Velichkovsky. Effective connectivity within the default mode network: dynamic causal modeling of resting-state fmri data. Frontiers in human neuroscience, 10:14, 2016.

[17] Michiel F Dirkx, Hanneke den Ouden, Esther Aarts, Monique Timmer, Bastiaan R Bloem, Ivan Toni, and Rick C Helmich. The cerebral network of parkinson’s tremor: an effective connectivity fmri study. Journal of Neuroscience, 36(19):5362–5372, 2016.

[18] Katrin H Preller, Adeel Razi, Peter Zeidman, Philipp Stämpfli, Karl J Friston, and Franz X Vollenweider. Effective connectivity changes in lsd-induced altered states of consciousness in humans. Proceedings of the National Academy of Sciences, 116(7):2743–2748, 2019.

[19] Edmund T Rolls, Wei Cheng, Matthieu Gilson, Jiang Qiu, Zicheng Hu, Hongtao Ruan, Yu Li, Chu-Chung Huang, Albert C Yang, Shih-Jen Tsai, et al. Effective connectivity in depression. Biological Psychiatry: Cognitive Neuroscience and Neuroimaging, 3(2):187–197, 2018.

[20] Elsie Premereur, Jessica Taubert, Peter Janssen, Rufin Vogels, and Wim Vanduffel. Effective connectivity reveals largely independent parallel networks of face and body patches. Current Biology, 26(24):3269–3279, 2016.

[21] Sebastian Gluth, Tobias Sommer, Jörg Rieskamp, and Christian Büchel. Effective connectivity between hippocampus and ventromedial prefrontal cortex controls preferential choices from memory. Neuron, 86(4):1078–1090, 2015.

[22] Vicente Pallarés, Andrea Insabato, Ana Sanjuán, Simone Kühn, Dante Mantini, Gustavo Deco, and Matthieu Gilson. Extracting orthogonal subject-and condition-specific signatures from fmri data using whole-brain effective connectivity. NeuroImage, 178:238–254, 2018.

[23] Scott L Fairhall and Alumit Ishai. Effective connectivity within the distributed cortical network for face perception. Cerebral cortex, 17(10):2400–2406, 2007.

[24] Tali Bitan, James R Booth, Janet Choy, Douglas D Burman, Darren R Gitelman, and M-Marsel Mesulam. Shifts of effective connectivity within a language network during rhyming and spelling. Journal of Neuroscience, 25(22):5397–5403, 2005.

[25] Christian Grefkes, Simon B Eickhoff, Dennis A Nowak, Manuel Dafotakis, and Gereon R Fink. Dynamic intra-and interhemispheric interactions during unilateral and bilateral hand movements assessed with fmri and dcm. Neuroimage, 41(4):1382–1394, 2008.

[26] Klaas E Stephan, Lee M Harrison, Stefan J Kiebel, Olivier David, Will D Penny, and Karl J Friston. Dynamic causal models of neural system dynamics: current state and future extensions. Journal of biosciences, 32(1):129–144, 2007.

[27] K.J. Friston, A. Mechelli, R. Turner, and C.J. Price. Nonlinear responses in fmri: The balloon model, volterra kernels, and other hemodynamics. NeuroImage, 12(4):466–477, 2000.

[28] Karl J Friston, Joshua Kahan, Bharat Biswal, and Adeel Razi. A dcm for resting state fmri. Neuroimage, 94:396–407, 2014.

[29] Adeel Razi, Joshua Kahan, Geraint Rees, and Karl J Friston. Construct validation of a dcm for resting state fmri. Neuroimage, 106:1–14, 2015.

[30] Giulia Prando, Mattia Zorzi, Alessandra Bertoldo, Maurizio Corbetta, Marco Zorzi, and Alessandro Chiuso. Sparse dcm for whole-brain effective connectivity from resting-state fmri data. NeuroImage, 208:116367, 2020.

[31] Giulia Vallini, Giorgia Baron, Erica Silvestri, Tommaso Volpi, AG Vlassenko, MS Goyal, Alessandro Chiuso, Diego Cecchin, Maurizio Corbetta, and Alessandra Bertoldo. Brain metabolic-functional (de) coupling from health to glioma dysfunction. Communications Biology, 2025.

[32] Stefan Frässle, Ekaterina I Lomakina, Adeel Razi, Karl J Friston, Joachim M Buhmann, and Klaas E Stephan. Regression dcm for fmri. NeuroImage, 155:406–421, 2017.

[33] Stefan Frässle, Samuel J. Harrison, Jakob Heinzle, Brett A. Clementz, Carol A. Tamminga, John A. Sweeney, Elliot S. Gershon, Matcheri S. Keshavan, Godfrey D. Pearlson, Albert Powers, and Klaas E. Stephan. Regression dynamic causal modeling for resting-state fmri. Human Brain Mapping, 42(7):2159–2180, 2021.

[34] Matthieu Gilson, Ruben Moreno-Bote, Adrián Ponce-Alvarez, Petra Ritter, and Gustavo Deco. Estimation of directed effective connectivity from fmri functional connectivity hints at asymmetries of cortical connectome. PLoS computational biology, 12(3):e1004762, 2016.

[35] David C Van Essen, Stephen M Smith, Deanna M Barch, Timothy EJ Behrens, Essa Yacoub, Kamil Ugurbil, Wu-Minn HCP Consortium, et al. The wu-minn human connectome project: an overview. Neuroimage, 80:62–79, 2013.

[36] Megan H Lee, Christopher D Smyser, and Joshua S Shimony. Resting-state fmri: a review of methods and clinical applications. American Journal of neuroradiology, 34(10):1866–1872, 2013.

[37] Jingyuan E Chen and Gary H Glover. Bold fractional contribution to restingstate functional connectivity above 0.1 hz. Neuroimage, 107:207–218, 2015.

[38] Danilo Benozzo, Giorgia Baron, Ludovico Coletta, Alessandro Chiuso, Alessandro Gozzi, and Alessandra Bertoldo. Macroscale coupling between structural and effective connectivity in the mouse brain. Scientific Reports, 14(1):3142, 2024.

[39] Giorgia Baron, Erica Silvestri, Danilo Benozzo, Alessandro Chiuso, and Alessandra Bertoldo. Revealing the spatial pattern of brain hemodynamic sensitivity to healthy aging through sparse dynamic causal model. Journal of Neuroscience, 45(1), 2025.

[40] Giorgia Baron, Giacomo Baggio, Massimiliano Facca, Danilo Benozzo, Alessandro Chiuso, Sandro Zampieri, and Alessandra Bertoldo. Eigenmode decomposition of asymmetries in whole-brain effective connectivity reveals multiscale hierarchical dynamics. bioRxiv, pages 2025–11, 2025.

[41] Matthieu Gilson, Gorka Zamora-López, Vicente Pallarés, Mohit H Adhikari, Mario Senden, Adrià Tauste Campo, Dante Mantini, Maurizio Corbetta, Gustavo Deco, and Andrea Insabato. Model-based whole-brain effective connectivity to study distributed cognition in health and disease. Network Neuroscience, 4(2):338–373, 2020.

[42] Matthieu Gilson, Nikos E Kouvaris, Gustavo Deco, Jean-François Mangin, Cyril Poupon, Sandrine Lefranc, Denis Rivière, and Gorka Zamora-López. Network analysis of whole-brain fmri dynamics: A new framework based on dynamic communicability. NeuroImage, 201:116007, 2019.

[43] Danilo Benozzo, Giacomo Baggio, Giorgia Baron, Alessandro Chiuso, Sandro Zampieri, and Alessandra Bertoldo. Analyzing asymmetry in brain hierarchies with a linear state-space model of resting-state fmri data. Network Neuroscience, 8(3):965–988, 10 2024.

[44] Enrico Amico and Joaquín Goñi. The quest for identifiability in human functional connectomes. Scientific reports, 8(1):8254, 2018.

[45] Dimitri Van De Ville, Younes Farouj, Maria Giulia Preti, Raphaël Liégeois, and Enrico Amico. When makes you unique: Temporality of the human brain fingerprint. Science advances, 7(42):eabj0751, 2021.

[46] Giovanni Donghi, Luca Pasa, Michele De Filippo De Grazia, Alberto Testolin, Marco Zorzi, Alessandro Sperduti, and Nicolò Navarin. On the application of neural networks for structured domains to fmri data. Neural Networks, 193:108029, 2026.

[47] Lucina Q Uddin, BT Thomas Yeo, and R Nathan Spreng. Towards a universal taxonomy of macro-scale functional human brain networks. Brain topography, 32(6):926–942, 2019.

[48] BT Thomas Yeo, Fenna M Krienen, Jorge Sepulcre, Mert R Sabuncu, Danial Lashkari, Marisa Hollinshead, Joshua L Roffman, Jordan W Smoller, Lilla Zöllei, Jonathan R Polimeni, et al. The organization of the human cerebral cortex estimated by intrinsic functional connectivity. Journal of Neurophysiology, 106(3):1125–1165, 2011.

[49] Tiago P Peixoto. Hierarchical block structures and high-resolution model selection in large networks. Physical Review X, 4(1):011047, 2014.

[50] Aaron F McDaid, Derek Greene, and Neil Hurley. Normalized mutual information to evaluate overlapping community finding algorithms. arXiv preprint 1110.2515, 2011.

[51] Sara Spadone, Stefania Della Penna, Carlo Sestieri, Viviana Betti, Annalisa Tosoni, Mauro Gianni Perrucci, Gian Luca Romani, and Maurizio Corbetta. Dynamic reorganization of human resting-state networks during visuospatial attention. Proceedings of the National Academy of Sciences, 112(26):8112–8117, 2015.

[52] Vinod Menon and Lucina Q Uddin. Saliency, switching, attention and control: a network model of insula function. Brain structure and function, 214(5):655– 667, 2010.

[53] Evan M Gordon, Charles J Lynch, Caterina Gratton, Timothy O Laumann, Adrian W Gilmore, Deanna J Greene, Mario Ortega, Annie L Nguyen, Bradley L Schlaggar, Steven E Petersen, et al. Three distinct sets of connector hubs integrate human brain function. Cell reports, 24(7):1687–1695, 2018.

[54] Xinjun Suo, Hao Ding, Xi Li, Yaodan Zhang, Meng Liang, Yongqiang Zhang, Chunshui Yu, and Wen Qin. Anatomical and functional coupling between the dorsal and ventral attention networks. Neuroimage, 232:117868, 2021.

[55] Dolly T Seeburger, Jason S Tsukahara, Nan Xu, Vishwadeep Ahluwalia, Shella D Keilholz, and Randall W Engle. Attention control ability is associated with frontoparietal control network interactions. Proceedings of the National Academy of Sciences, 123(19):e2526828123, 2026.

[56] Manesh Girn, Roni Setton, Gary R Turner, and R Nathan Spreng. The “limbic network,” comprising orbitofrontal and anterior temporal cortex, is part of an extended default network: Evidence from multi-echo fmri. Network Neuroscience, 8(3):860–882, 2024.

[57] Yuan Zhou, Karl J Friston, Peter Zeidman, Jie Chen, Shu Li, and Adeel Razi. The hierarchical organization of the default, dorsal attention and salience networks in adolescents and young adults. Cerebral cortex, 28(2):726–737, 2018.

[58] Hideya Koshino, Mariko Osaka, Tetsuya Shimokawa, Mizuki Kaneda, Seira Taniguchi, Takehiro Minamoto, Ken Yaoi, Miyuki Azuma, Katsuki Higo, and Naoyuki Osaka. Cooperation and competition between the default mode network and frontal parietal network in the elderly. Frontiers in psychology, 14:1140399, 2023.

[59] Jarod L Roland, Abraham Z Snyder, Carl D Hacker, Anish Mitra, Joshua S Shimony, David D Limbrick, Marcus E Raichle, Matthew D Smyth, and Eric C Leuthardt. On the role of the corpus callosum in interhemispheric functional connectivity in humans. Proceedings of the National Academy of Sciences, 114(50):13278–13283, 2017.

[60] Lenny E Ramsey, Joshua S Siegel, Antonello Baldassarre, Nicholas V Metcalf, Kristina Zinn, Gordon L Shulman, and Maurizio Corbetta. Normalization of network connectivity in hemispatial neglect recovery. Annals of neurology, 80(1):127–141, 2016.

[61] Mohit H Adhikari, Joseph Griffis, Joshua S Siegel, Michel Thiebaut de Schotten, Gustavo Deco, Andrea Instabato, Matthieu Gilson, and Maurizio Corbetta. Effective connectivity extracts clinically relevant prognostic information from resting state activity in stroke. Brain communications, 3(4):fcab233, 2021.

[62] Matthieu Gilson, Enzo Tagliazucchi, and Rodrigo Cofré. Entropy production of multivariate ornstein-uhlenbeck processes correlates with consciousness levels in the human brain. Physical Review E, 107(2):024121, 2023.

[63] D Rangaprakash, Guo-Rong Wu, Daniele Marinazzo, Xiaoping Hu, and Gopikrishna Deshpande. Hemodynamic response function (hrf) variability confounds resting-state fmri functional connectivity. Magnetic resonance in medicine, 80(4):1697–1713, 2018.

[64] Guo-Rong Wu, Wei Liao, Sebastiano Stramaglia, Ju-Rong Ding, Huafu Chen, and Daniele Marinazzo. A blind deconvolution approach to recover effective connectivity brain networks from resting state fmri data. Medical image analysis, 17(3):365–374, 2013.

[65] Christian Lohse, Danielle S Bassett, Kelvin O Lim, and Jean M Carlson. Resolving anatomical and functional structure in human brain organization: identifying mesoscale organization in weighted network representations. PLoS computational biology, 10(10):e1003712, 2014.

[66] Victor Buendía, Pablo Villegas, Raffaella Burioni, and Miguel A Muñoz. The broad edge of synchronization: Griffiths effects and collective phenomena in brain networks. Philosophical Transactions of the Royal Society A, 380(2227):20200424, 2022.

[67] Giacomo Barzon, Oriol Artime, Samir Suweis, and Manlio De Domenico. Unraveling the mesoscale organization induced by network-driven processes. Proceedings of the National Academy of Sciences, 121(28):e2317608121, 2024.

[68] Richard F Betzel and Danielle S Bassett. Multi-scale brain networks. Neuroimage, 160:73–83, 2017.

[69] Markus Axer and Katrin Amunts. Scale matters: The nested human connectome. Science, 378(6619):500–504, 2022.

[70] Ana Clara Caznok Silveira, Andre Saraiva Leao Marcelo Antunes, Maria Carolina Pedro Athié, Bárbara Filomena da Silva, João Victor Ribeiro dos Santos, Camila Canateli, Marina Alves Fontoura, Allan Pinto, Luciana Ramalho Pimentel-Silva, Simoni Helena Avansini, et al. Between neurons and networks: investigating mesoscale brain connectivity in neurological and psychiatric disorders. Frontiers in neuroscience, 18:1340345, 2024.

[71] Jivesh Ramduny and Clare Kelly. Connectome-based fingerprinting: reproducibility, precision, and behavioral prediction. Neuropsychopharmacology, 50(1):114–123, 2025.

[72] Pierpaolo Sorrentino, Rosaria Rucco, Anna Lardone, Marianna Liparoti, Emahnuel Troisi Lopez, Carlo Cavaliere, Andrea Soricelli, Viktor Jirsa, Giuseppe Sorrentino, and Enrico Amico. Clinical connectome fingerprints of cognitive decline. NeuroImage, 238:118253, 2021.

[73] Sara Stampacchia, Saina Asadi, Szymon Tomczyk, Federica Ribaldi, Max Scheffler, Karl-Olof Lövblad, Michela Pievani, Aïda B Fall, Maria Giulia Preti, Paul G Unschuld, et al. Fingerprints of brain disease: connectome identifiability in alzheimer’s disease. Communications Biology, 7(1):1169, 2024.

[74] Sara Stampacchia, Fosco Bernasconi, Dimitri Van De Ville, Enrico Amico, Javier Pagonabarraga, Jaime Kulisevsky, and Olaf Blanke. Connectome-based brain fingerprints predict early cognitive decline in parkinson’s patients with minor hallucinations. bioRxiv, pages 2025–04, 2025.

[75] Tobias Kaufmann, Dag Alnæs, Christine L Brandt, Francesco Bettella, Srdjan Djurovic, Ole A Andreassen, and Lars T Westlye. Stability of the brain functional connectome fingerprint in individuals with schizophrenia. JAMA psychiatry, 75(7):749–751, 2018.

[76] Lorenzo Cipriano, Emahnuel Troisi Lopez, Marianna Liparoti, Roberta Minino, Antonella Romano, Arianna Polverino, Francesco Ciaramella, Michele Ambrosanio, Simona Bonavita, Viktor Jirsa, et al. Reduced clinical connectome fingerprinting in multiple sclerosis predicts fatigue severity. NeuroImage: Clinical, 39:103464, 2023.

[77] Gustavo Deco, Diego Vidaurre, and Morten L Kringelbach. Revisiting the global workspace orchestrating the hierarchical organization of the human brain. Nature human behaviour, 5(4):497–511, 2021.

[78] Karan Kabbur Hanumanthappa Manjunatha, Giorgia Baron, Danilo Benozzo, Erica Silvestri, Maurizio Corbetta, Alessandro Chiuso, Alessandra Bertoldo, Samir Suweis, and Michele Allegra. Controlling target brain regions by optimal selection of input nodes. PLOS Computational Biology, 20(1):1–30, 01 2024.

[79] Jun Cao, Yifan Zhao, Xiaocai Shan, Hua-liang Wei, Yuzhu Guo, Liangyu Chen, John Ahmet Erkoyuncu, and Ptolemaios Georgios Sarrigiannis. Brain functional and effective connectivity based on electroencephalography recordings: A review. Human brain mapping, 43(2):860–879, 2022.

[80] Zixiang Luo, Kaining Peng, Zhichao Liang, Shengyuan Cai, Chenyu Xu, Dan Li, Yu Hu, Changsong Zhou, and Quanying Liu. Mapping effective connectivity by virtually perturbing a surrogate brain. Nature Methods, 22(6):1376–1385, 2025.

[81] Leonardo Novelli, Karl Friston, and Adeel Razi. Spectral dynamic causal modeling: A didactic introduction and its relationship with functional connectivity. Network Neuroscience, 8(1):178–202, 2024.

[82] Matthew D Greaves, Leonardo Novelli, Sina Mansour L, Andrew Zalesky, and Adeel Razi. Structurally informed models of directed brain connectivity. Nature Reviews Neuroscience, 26(1):23–41, 2025.

[83] Francesca Mastrogiuseppe, Joana Carmona, M Yu Byron, Adam Kohn, and Christian K Machens. Input-dependent directionality of interactions between cortical areas. bioRxiv, 2026.

[84] Davide Borra and Elisa Magosso. A deep learning-enriched framework for analyzing brain functional connectivity. Scientific Reports, 15(1):34504, 2025.

[85] Peter Zeidman, Amirhossein Jafarian, Nadège Corbin, Mohamed L Seghier, Adeel Razi, Cathy J Price, and Karl J Friston. A guide to group effective connectivity analysis, part 1: First level analysis with dcm for fmri. Neuroimage, 200:174–190, 2019.

[86] Matthew F Glasser, Stamatios N Sotiropoulos, J Anthony Wilson, Timothy S Coalson, Bruce Fischl, Jesper L Andersson, Junqian Xu, Saad Jbabdi, Matthew Webster, Jonathan R Polimeni, et al. The minimal preprocessing pipelines for the human connectome project. Neuroimage, 80:105–124, 2013.

[87] Alexander Schaefer, Ru Kong, Evan M Gordon, Timothy O Laumann, Xi-Nian Zuo, Avram J Holmes, Simon B Eickhoff, and BT Thomas Yeo. Local-global parcellation of the human cerebral cortex from intrinsic functional connectivity mri. Cerebral cortex, 28(9):3095–3114, 2018.

[88] Yusi Chen, Burke Q Rosen, and Terrence J Sejnowski. Dynamical differential covariance recovers directional network structure in multiscale neural systems. Proceedings of the National Academy of Sciences, 119(24):e2117234119, 2022.

[89] Tiger W Lin, Anup Das, Giri P Krishnan, Maxim Bazhenov, and Terrence J Sejnowski. Differential covariance: A new class of methods to estimate sparse connectivity from neural recordings. Neural computation, 29(10):2581–2632, 2017.

[90] Roger Guimera, Marta Sales-Pardo, and Luís A Nunes Amaral. Modularity from fluctuations in random graphs and complex networks. Physical Review E—Statistical, Nonlinear, and Soft Matter Physics, 70(2):025101, 2004.

[91] Tiago P. Peixoto. Revealing consensus and dissensus between network partitions. Phys. Rev. X, 11:021003, Apr 2021.

